# CD5L insufficiency exacerbates skeletal joint damage in rheumatoid arthritis

**DOI:** 10.64898/2025.12.16.694616

**Authors:** Diana Bicho, Malin C. Erlandsson, David Svensson, Emma Comas, Maria Carolina Silva, Venkataragavan Chandrasekaran, Victoria M.E. Lundgren, Rita F. Santos, Liliana Oliveira, Maria I. Bokarewa, Alexandre M. Carmo

## Abstract

CD5L is an immunoregulatory protein induced during inflammation and makes a life-saving contribution in infection and sepsis. Here, we explored the overall impact of CD5L in rheumatoid arthritis (RA). Using experimental RA models, we demonstrated an early surge of CD5L expression in wild-type mice, as well as a higher incidence and increased severity of arthritis in CD5L-deficient mice. In the blood, CD5L-deficient mice exhibited enhanced inflammation, a higher proportion of CD11b⁺ mononuclear cells, and elevated levels of IL-1β, IL-17, and IFN-γ, particularly at the pre-clinical stage of arthritis. In human RA, mononuclear cells conditioned with CD5L, as well as endogenous CD5L production, were associated with reduced CD11b expression, increased IL-10 and PD-L1 production, and an enrichment of non-classical CD16^+^ monocytes. Analysis of the CD5L-dependent transcriptome in CD14⁺ cells from RA patients revealed the acquisition of an efferocytosis profile, with upregulation of C1Q subunits, GAS6, AXL, and ALOX15B. It also showed signs of insufficient cargo processing, which dampens the resolution of inflammation, permits the expansion of IFN-primed non-classical monocytes, and correlates strongly with joint damage accrual. Despite markedly low CD5L levels in the synovium, RA synovial tissue was enriched with non-classical monocytes carrying the GAS6-AXL signature, which fosters osteoclast progenitors. In conclusion, this study demonstrates that the anti-inflammatory and inflammation-resolving properties of CD5L in experimental and human RA are mediated through the induction of efferocytosis-related and IFN-primed monocyte programs. This harbors a potential risk of uncontrolled invasion of non-classical monocytes into the synovial tissue, leading to joint structural damage. The complex role of CD5L in inflammation and disease outcomes requires careful consideration of the disease phase and experimental context.

## 1. Introduction

CD5 antigen-like (CD5L) is a circulating glycoprotein abundant in blood, best known for its ability to inhibit hematopoietic cell apoptosis [1, 2]. It is primarily produced by monocytes and macrophages in the spleen, lymphoid tissues, and bone marrow [3], and, among other immunoregulatory roles, recognizes and neutralizes pathogen-associated molecular patterns [4, 5], and damage-associated molecular patterns released from dying host cells, facilitating debris clearance and limiting tissue inflammation [6–8].

Circulating CD5L levels are elevated in several infectious and inflammatory conditions, yet whether it acts protectively or pathogenically remains uncertain, particularly in diseases where its role is poorly characterized [9]. Insights into CD5L function have largely come from studies using CD5L-deficient mice, which reveal diverse outcomes depending on the disease model. In experimental sepsis models, CD5L-deficient mice show impaired neutrophil recruitment, reduced bacterial control, and higher mortality in polymicrobial cecal ligation and puncture (CLP) and LPS-induced septic shock models [10]. CD5L also restrains adaptive immune pathogenicity; it limits the pathogenic program of Th17 cells, and its absence shifts Th17 cells toward a more pathogenic phenotype [11]. In contrast, CD5L deletion is protective in several non-infectious injury models. CD5L-deficient mice are less susceptible to bleomycin-induced pulmonary fibrosis [12] and to acetaminophen-induced acute liver injury [13]. In yet other contexts, such as *Mycobacterium tuberculosis* infection, CD5L deficiency has no detectable impact on overall disease outcome, even though CD5L levels rise upon infection [14]. Together, these studies demonstrate that CD5L can be protective, pathogenic, or neutral depending on the model and tissue context.

In humans, increased serum CD5L has been reported in several conditions, including atopic dermatitis [15], systemic lupus erythematosus [16], liver fibrosis [17], Alzheimer’s disease [18], and Kawasaki disease [19]. Recent evidence also suggests that CD5L may be relevant in autoimmune joint disease: in a limited patient cohort, circulating CD5L levels were elevated in rheumatoid arthritis (RA) and correlated with disease severity [20]. Although this association identifies CD5L as a potential biomarker of inflammation in RA, its functional contribution to disease pathogenesis remains poorly understood.

RA is an inflammatory joint disease with an estimated global prevalence of 0.5-1.0% [21]. Morphologically, it is characterized by leukocyte infiltration into the joint tissue, synovial hyperplasia, and progressive skeletal damage [22]. Monocytes and macrophages play a pivotal role in RA pathogenesis by driving leukocyte infiltration, maintaining a pro-inflammatory cytokine milieu, and inducing synovial hyperplasia and bone remodeling [23, 24].

Traditionally, the surface markers CD14 and FCGR3A (CD16a) classify circulating monocytes into three major subsets: classical (CD14⁺CD16⁻), intermediate (CD14⁺CD16⁺), and non-classical (CD14^low^CD16⁺) [25]. Functionally, classical monocytes are highly phagocytic and pro-inflammatory, intermediate monocytes produce high levels of inflammatory cytokines and contribute to antigen presentation, while non-classical monocytes patrol the endothelium and support tissue repair [26]. In RA, monocytes are recruited from the peripheral blood to synovial tissue via chemokines such as CCL2 and CX3CL1 [27, 28], and within the synovial environment, signals from cytokines (IL-6, IL-1β, TNF-α, and IFN-γ), fibroblasts, hypoxia, and immune complexes engaging Fcγ receptors drive their differentiation into joint-resident macrophages [29–34].

Single-cell analyses of RA synovial biopsies have further refined the identification of distinct macrophage subsets with specialized roles in disease pathogenesis. One study identified a pro-inflammatory population characterized by high production of IL-6 and TNF and lacking expression of the receptor tyrosine kinase MerTK, a key molecule involved in efferocytosis [35]. Another study revealed a distinct subset of inflammatory macrophages that promotes fibroblast aggressiveness and invasion through expression of heparin-binding EGF-like growth factor (HB-EGF), urokinase receptor (UPAR), and NR4A family transcription factors [36]. In contrast, CX3CR1⁺MerTK⁺ macrophages function as organizers of the immunological barrier and support resolution of inflammation [37–39].

The emergence and maintenance of macrophage phenotypic states are strongly influenced by epigenetic mechanisms. In RA, an altered balance between histone acetyltransferase (HAT) and histone deacetylase (HDAC) activities leads to global histone hyperacetylation, increasing chromatin accessibility at both inflammatory and regulatory gene loci [40, 41]. Reduced HDAC activity, particularly HDAC3, contributes to this hyperacetylation, amplifying NF-κB-driven IL-6 and TNF production [42, 43]. Conversely, certain HDACs, including SIRT1, can epigenetically promote IL-10 expression, driving macrophage polarization toward a pro-resolving M2 phenotype [41]. Thus, histone acetylation acts as a context-dependent regulatory mechanism capable of either sustaining inflammation or facilitating its resolution.

Despite advances in characterizing the cellular and epigenetic drivers of RA, the contribution of CD5L to disease development remains poorly explored. In this study, we investigated the role of CD5L in RA pathogenesis using the collagen-induced arthritis (CIA) model and profiling the inflammatory response in CD5L-deficient mice throughout disease progression. Guided by these experimental findings, we then assessed clinical associations of serum CD5L levels in RA patients and characterized the monocyte phenotypes linked to CD5L production. Overall, our results indicate that CD5L exerts protective anti-inflammatory effects in RA, although its endogenous activity seems insufficient to fully prevent skeletal damage in affected joints.

## 2. Materials and methods

### 2.1. Ethics statement

All animal experiments were conducted in strict accordance with the Portuguese (Portaria 1005/92) and European (Directive 2010/63/EU) legislations governing the housing, husbandry, and welfare of laboratory animals. The study protocols were reviewed and approved by the Ethics Committee of the Instituto de Investigação e Inovação em Saúde (i3S), Universidade do Porto, and by the Portuguese National Entity Direção Geral de Alimentação e Veterinária (license references: 022868/2020-12-31 and 009951/2018-05-17).

Human studies received the approval from the board of the Ethical Committee of Gothenburg, Sweden (Cohort 1: Dnr 659-11; Cohort 2: Dnr 2019-03787). The collection of biological material was conducted in accordance with national regulations and the International Conference on Harmonization Good Clinical Practice requirements, based on the Declaration of Helsinki.

### 2.2. Mice and experimental procedures

CD5L knockout (*Cd5l*^−/−^, hereafter referred to as CD5L-deficient or CD5L⁻) mice on a C57BL/6J background, previously generated and characterized in our laboratory [10], were maintained under specific pathogen-free conditions at the i3S animal facility (Porto, Portugal). Wild-type (*Cd5l*^+/+^, hereafter WT) littermates were used as controls. Peripheral blood analyses revealed no major differences in leukocyte counts or subset distribution between WT and CD5L⁻ mice [10].

All experiments were performed using 8–12-week-old male mice bred in-house. Male mice were selected due to their higher susceptibility to CIA. Animals were housed at 21 ± 2 °C under a 12-h light/dark cycle, with ad libitum access to food and water, in an AAALAC-accredited facility.

### 2.3. Induction and evaluation of collagen-induced arthritis (CIA)

#### Induction of CIA

Arthritis was induced as previously described [44], replacing bovine with chicken type II collagen (CII) (Chondrex, cat. no. 20012, Woodinville, WA, USA). On day 0, mice were immunized subcutaneously at the base of the tail with 0.2 mg of chicken CII emulsified in an equal volume of complete Freund’s adjuvant (CFA, Chondrex, cat. no. 7023; 0.5 mg/mouse; total volume 200 μl). On day 21, mice received a booster injection of CII in incomplete Freund’s adjuvant (Sigma-Aldrich, cat. no. F5506, St. Louis, MO, USA).

#### Arthritis evaluation

Arthritis severity was evaluated in all four paws using a blinded scoring system by a trained observer. Each paw was scored for swelling and erythema: 0 = normal, 1 = mild swelling/erythema, 2 = pronounced swelling, 3 = deformity, 4= ankylosis. The maximum score achieved for each joint (ankle, tarsus, carpus, toes and finger) was 4, so the maximum arthritis score for a single mouse on a given day could reach 40. Mice were evaluated two to three times per week, and body weight was recorded weekly.

#### Blood collection

Blood samples were drawn using the facial vein. For that, animals were anesthetized with inhaled isoflurane, blood was collected at the different time points using a 23G needle followed by aspiration into a 75 μl heparinized microhematocrit tube (Fisher Scientific, Hampton, NH, USA), and stored in Eppendorf tubes for 30 min at RT. Subsequently, the serum was separated from the clotted blood by centrifugation at 9,000 × *g* at 4 °C for 10 min and stored at –20 °C until further use.

#### Immunophenotyping of mouse leukocytes

Blood was treated with red blood cell lysis buffer (0.01 M Tris, 0.15 M NH_4_Cl, pH 7.2) at RT for 5 min. Subsequently, the cells were centrifuged at 300 × *g* for 5 min at 4 °C, washed in 50 μl FACS buffer [0.2% (w/v) BSA + 0.1% (w/v) NaN_3_ in PBS] and centrifuged again and blocked using FcR-block [TruStain FcX™ (anti-mouse CD16/32), BioLegend, San Diego, CA, USA] for 15 min on ice. After a new centrifugation, the cells were incubated for 30 min with the following antibodies (all BioLegend, San Diego, CA, USA unless otherwise stated): CD3 APC-Cy7 (17A2), CD19 APC (6D5), NK1.1 FITC (PK136), CD11b PE (M1/70), F4/80 APC-Cy7 (BM8), Ly6C FiTC (HK1.4), Ly6G PB (1A8), and Siglec-F Alexa Fluor® 647 (E50-2440, BD Biosciences, Franklin Lakes, NJ, USA). After washes, cells were resuspended in FACS buffer, filtered using a 0.2 μm cell strainer and data were acquired on a three-laser BD FACSCanto II (Becton Dickinson). Data analysis was performed in FlowJo v10.10.0 (BD Biosciences).

The phenotyping gating strategy is shown in Fig. S1. Cells were identified as T cells (CD3⁺), B cells (CD19⁺), NK cells (NK1.1⁺CD3⁻), eosinophils (SSC^hi^Siglec-F⁺CD11b⁺), neutrophils (Siglec-F⁻Ly6G⁺CD11b⁺), monocytes (Siglec-F⁻Ly6G⁻F4/80⁺CD11b⁺), classical/inflammatory monocytes (Siglec-F⁻Ly6G⁻F4/80⁺CD11b⁺Ly6C⁺), and patrolling/non-classical monocytes (Siglec-F⁻Ly6G⁻ F4/80⁺ CD11b⁺ Ly6C⁻).

#### Quantification of soluble factors

Bone resorption markers RANKL and cross-linked C-telopeptide of type I collagen (CTX-I) were measured in the sera of WT and CD5L^−^ mice using the Mouse RANKL (soluble) ELISA development Kit (#900-K233K, Peprotech) and Mouse C-telopeptide of type I collagen, CTX-I ELISA Kit (#MBS164705, MybioSource), respectively, according to the manufacturers’ instructions. Cytokine levels in the serum of WT and CD5L^−^ mice were measured using ELISA Max™ Deluxe Sets (BioLegend) for IL-17A (#432504), IL-1β (#437004), IL-6 (#431304), and IFN-γ (#430804). CD5L serum levels were quantified using the Mouse CD5L ELISA Pair Set (Sino Biological, SEK50020), according to the manufacturers’ instructions.

Anti-CII antibodies were quantified as follows: microtiter plates (Nunc MaxiSorp™, 44-2404-21, Thermo Fisher Scientific) were coated with 5 μg/ml CII dissolved in coating buffer (0.05 M Tris, 0.2 M NaCl, pH 7.4) overnight at 4 °C. The plates were washed with 0.05 % (v/v) Tween 20 in PBS and blocked with 5% fetal bovine serum in PBS for 1 h. Then, the test serum was serially diluted and plated, interacting with the coated collagen for 2 h. Total bound IgG was detected with goat anti-mouse IgG-HRP (Poly4053, BioLegend), followed by substrate solution (SIGMAFAST™ OPD, P9187, Sigma-Aldrich). Reactions were stopped with 3 N H_2_SO_4_ and the optical density was measured at 492 nm using a μQuant™ Microplate Reader (BioTek Instruments). Results for each sample were graphed using GraphPad Prism, with the 1/serum dilution on the x-axis (log scale) versus the optical density (OD) on the y-axis (linear scale). In the region where the line on the graph was linear for all samples, the lowest possible OD that intersected all of the samples in that region was chosen and the corresponding titer for each sample was obtained from the graph, with adjustment for specific IgG isotype and allotype sensitivity, as described elsewhere [45]. A pool of sera collected from CII-immunized mice was similarly diluted to that of the test samples and included on each plate. This serum pool was used as standard to assure reproducibility and was adjusted for comparison of results from plate to plate.

### 2.4. Human samples

#### Cohorts and clinical data

In total, the study included a cross-sectional cohort of 80 female patients with RA (mean age 60 ± 12 years; mean disease duration 14.7 years) who were attending the Rheumatology Clinic at Sahlgrenska University Hospital, Gothenburg, for routine follow-up visits. Cohort 1 was collected during the period 2012-2013, cohort 2 was collected during 2019. Demographic and clinical characteristics of the patient cohorts are summarized in Table 1. In 2016, CD5L was measured in the serum of 40 patients with RA, 11 healthy controls, and an additional 5 paired samples of blood and synovial fluid. In 2024, we measured CD5L levels in 35 RA patients and in supernatants of CD14⁺ cells isolated from the same individuals.

**Table 1.** Clinical characteristics of the patients with rheumatoid arthritis.

| all female | Cohort 1+2<br>n=80 | Cohort 2 (RNAseq)<br>n=35 |
| --- | --- | --- |
| Age, y mean (min-max) | 60.9 (29-76) | 64.9 (46-76) |
| DD, y mean (min-max) | 14.7 (1-45) | 12.8 (1-45) |
| AB-positive | 69 (86.2) | 28 (80%) |
| RF-positive | 66 (82.5) | 26 (74.3%) |
| ACPA-positive | 62 (77.5) | 18 (51.4%) |
| DAS28, mean (min-max) | 3.17 (1.07-5.77) | 2.87 (1.07-5.77) |
| Total Sharp-vdH score, mean (min-max) | 52.0 (0-299) | 44.8 (8-58) |
| Treatment |  |  |
| MTX | 64 (80%) | 24 (68.6%) |
| TNF-inhibitors | 51 (63.6%) | 10 (28.6%) |
| Other DMARDs | 5 (6.2%) | 2 (5.7%) |

DD, disease duration; AB, antibodies; RF, rheumatoid factor; ACPA, antibodies to citrullinated peptides; DAS28 disease activity score by 28 joints; Sharp-vdH, Sharp van der Heide; MTX, methotrexate; TNF, tumor necrosis factor; DMARDs, disease modifying anti-rheumatic drugs.

Disease-modifying antirheumatic drugs (DMARDs) were used by 73 of the 80 patients, including methotrexate (MTX) in 64 patients. Five patients used other drugs (azathioprine, n = 3; chlorambucil, n = 1; leflunomide, n = 1). TNF inhibitors were administered to 51 patients, all in combination with MTX or another drugs. All patients and healthy controls provided written informed consent prior to participation in any study-related procedures.

#### Sample collection and processing

Blood samples were collected in sterile vacuum tubes containing sodium citrate (0.129 mol/l, pH 7.4) for plasma, and in additive-free tubes for serum. Samples were then placed on ice until centrifugation (1600 × *g*, 4 °C, 15 min). Serum and plasma supernatants were aliquoted into RNAse/DNAse-free tubes and stored at –80 °C until further use.

Synovial fluid samples were obtained by arthrocentesis of knee joints prior to routine corticosteroid injection. SF was aspirated aseptically and transferred into tubes containing sodium citrate, aliquoted, and stored at –80 °C until analysis. All samples were coded and analyzed by investigators blinded to the clinical and radiological data.

#### Clinical data collection

A standard protocol for evaluation of joints was used considering the swelling, tenderness, limited motion or deformity of each joint [46]. Disease activity was measured with the disease activity score in 28 joints (DAS28), including 10 metacarpal phalangeal and 10 proximal interphalangeal joints of the hands, 2 shoulders, 2 elbows, 2 wrists, and 2 knees. The DAS28 scores were calculated based on the number of tender and swollen joints, and erythrocyte sedimentation rate (ESR) [47].

#### Radiographic evaluation

The degree of radiographic joint damage was calculated from posterior-anterior radiographs of the hands and feet using the Sharp method modified by van der Heijde (vdH-Sharp) [48]. The vdH-Sharp score was presented as a total score (range: 0–448), an erosion score (0–280), and a joint space narrowing score (0–168). The total vdH-Sharp > 50 characterized severe radiographic damage.

### 2.5. Serological measurements

CD5L levels in serum and synovial fluid were quantified using the human CD5L ELISA kit (ab213760; Abcam, Cambridge, UK); detection limit, 10 pg/ml. CD5L levels from serum from RNAseq samples and supernatants from PBMC and CD14^+^ cell cultures were quantified using the human CD5L ELISA kit (EH94RB, Invitrogen, Waltham, MA, USA); detection limit, 0.3 pg/ml.

Cartilage oligomeric matrix protein (COMP) levels were measured by an ELISA kit (Anamar, Lund, Sweden) in samples diluted 1:10; detection limit, 0.9 IU/ml. C-telopeptide cross-link type I collagen (CTX-I) was measured by an ELISA (IDS Serum CrossLaps, Immunodiagnostic Systems, Boldon, UK) in undiluted samples; detection limit, 0.11 ng/ml. Matrix metalloproteinase 3 (MMP-3) levels were measured by an ELISA using a pair of matched antibodies (DY513, R&D Systems, Minneapolis, MN, USA) in samples diluted 1:100; detection limit, 3 ng/ml. Serum S100A4 levels were measured by an ELISA (KA0092, Abnova, Taipei City, Taiwan) in samples diluted 1:200; detection limit, 10 ng/ml.

Cytokine levels were also determined by ELISA: IL-10 (DY217B, R&D Systems,; detection limit, 3 pg/ml), IFN-γ (DY285B, R&D Systems; detection limit, 5 pg/ml), and PD-L1 (DY156, R&D Systems; detection limit, 0.5 pg/ml). Erythrocyte sedimentation rate (ESR) was measured by the Westergren method, and C-reactive protein (CRP) was determined by nephelometry at the hospital laboratory.

In cohort 1, IL-6 levels were measured by a bioassay using the murine hybridoma cell line B9, which is dependent on IL-6 for growth [49]; sample dilution, 1:50; detection limit, 50 pg/ml. In cohort 2, IL-6 levels were analyzed by ELISA (Sanquin, Amsterdam, The Netherlands).

In cohort 1, IGF-1 levels were measured by an ELISA using a pair of matched antibodies (DY291, R&D Systems) in samples diluted 1:25; detection limit, 1 ng/ml. In cohort 2, IGF-1 levels were analyzed by photometry on the Cobas 8000 system (Roche Diagnostics, Basel, Switzerland). All detection limits are reported according to the manufacturers’ instructions.

### 2.6. Isolation and stimulation of human CD14^+^ cells

Human peripheral blood mononuclear cells (PBMCs) were isolated from heparinized venous peripheral blood by density gradient separation on Lymphoprep (Axis-Shield PoC AS, Dundee, Scotland). PBMC cultures (1 × 10⁶ cells/ml) were propagated in RPMI medium (Gibco, Waltham, MA, USA) containing 50 μM β-mercaptoethanol (Gibco), 2 mM GlutaMAX (Gibco), 50 μg/ml gentamicin (Sanofi-Aventis, Paris, France), and 5% fetal bovine serum (Sigma-Aldrich) at 37 °C in a humidified 5% CO₂ atmosphere.

Cells were stimulated with concanavalin A (ConA, 0.625 μg/ml; Sigma-Aldrich) and lipopolysaccharide (LPS, serotype O55:B5, 5 μg/ml; Sigma-Aldrich) for 48 h. After 24 h, some cultures were supplemented with HDAC inhibitor (HDACi) valproate (50 μg/ml; Ergenyl, Sanofi, Paris, France), human recombinant CD5L (1 μg/ml (INVIGATE GmbH, Jena, Germany), human IgG (4 μg/ml; Kiovig, Takeda, Tokyo, Japan), or a combination of CD5L with human IgG, as indicated. The synthetic Lyn activator talimogene (1 and 10 μg/ml; Selleckchem, Houston, USA) and the proteasome inhibitor bortezomib (500 ng/ml; Stada Nordic, Herlev, Denmark) were also used.

CD14⁺ cells were isolated from fresh PBMC cultures by positive selection (kit #17858; StemCell Technologies, Vancouver, Canada) and propagated in the enriched RPMI medium described above at a concentration of 1.25 × 10⁶ cells/ml. For RNAseq and qPCR, cells were stimulated with LPS (5 μg/ml) for 2 h.

At the end of incubation, supernatants were collected for cytokine measurement, and cells were analyzed by flow cytometry. Adherent cells were detached by incubation with 1 mM ice-cold EDTA for 15 min, followed by gentle mechanical scraping. Adherent and non-adherent cells were then pooled for subsequent analysis.

### 2.7. Flow cytometry

Cells were resuspended in PBS containing 2% fetal calf serum, 0.1% NaN_3_ and 1 mM EDTA (FACS buffer). Non-specific binding was blocked with human γ-globulin (100 μg/ml, Kiovig, Takeda), thereafter monoclonal mouse anti-human antibodies binding CD4-Brilliant Violet 510 (Clone SK3, Beckton Dickinson (BD), Franklin Lakes, NJ, USA), CD14-PE (Clone M5E2, BD), CD16-FITC (Clone ME5E2, BD), CD29-APC (Clone MAR4, BD), CD11b-Pacific Blue (Clone CRF44, BD), CD45-PerCP (Clone 2D1, BD) were added. All antibodies were diluted in FACS buffer to optimal concentrations.

Cells (100,000 events/sample) were acquired on a FACS-Lyric (Beckton Dickinson) and analyzed with FlowJo v10.10 (Beckton Dickinson). Cell populations were gated using fluorochrome minus one (FMO) staining controls. The total monocyte population was analyzed within the CD4^−^CD11b^+^ gate. Within this, three major monocyte subsets were identified: CD14^+^CD16^−^ (classical monocytes); CD14^+^CD16^+^ (intermediate monocytes); and CD14^low^CD16^+^ (non-classical monocytes). Data are presented as the percentage of CD4^−^CD11b^+^ cells or as mean fluorescence intensity (MFI) in positive cells.

### 2.8. Transcriptome sequencing (RNA-seq) and analysis

Total RNA from CD14^+^ cells was prepared using the Norgen Total RNA kit (17200 Norgen Biotek, Ontario, Canada). Quality control was done by Bioanalyzer RNA6000 Pico on Agilent2100 (Agilent, Santa Clara, CA, USA). Deep sequencing was done by RNAseq (Hiseq2000, Illumina) at the core facility for Bioinformatics and Expression Analysis (Karolinska Institute, Huddinge, Sweden). Raw sequence data were obtained in Bcl-files and converted into fastq text format using the bcl2fastq program from Illumina.

Analysis was performed using the core Bioconductor packages in R-studio v. 4.4.1. Mapping of transcripts was done using Genome UCSC annotation set for hg38 human genome assembly. Using the normalized expression data in CD14^+^ cells of 35 RA patients, the ‘rank metric score’ was used to rank the protein coding genes. For each gene transcript, the rank metric score was calculated as difference of the mean expression scaled by the standard deviation and identified the genes acting as reliable markers of this gene set. The genes with the rank metric score above 0.200 were used in the linear regression analysis of z-standardized serum CD5L levels and of CD5L levels in supernatants of CD14^+^ cells to the transcriptome.

Using a design formula as ∼log_CD5L_scaled in the DESeq2 pipeline, we identified protein-coding genes with a base mean expression ≥ 10 that exhibited a significant change (|log₂FC| > 0, nominal *p* ≤ 0.05) per unit increase in serum CD5L level. These genes were subjected to the analysis for the enriched biological pathways through the Gene Set Enrichment Analysis (GSEA) online tool from Broad Institute, UC San Diego, USA.

#### Analysis of Single cell transcriptome in myeloid cells of synovial tissue

To analyze CD5L-dependent gene signatures, we used the single cell transcriptome dataset of synovial tissue of RA patients [50]. The log-normalized mRNA matrix was used to create the Seurat object and subsetted to the myeloid cell cluster (n=76,181 cells). Within this subset, the joint density of gene expression for different gene signatures was calculated and visualized using the do_NebulosaPlot function from the SCpubr package [51]. Dotplots of CD5L expression were visualized using the Dotplot_scCustom function from the scCustomize package [52].

### 2.9. Data availability

Transcriptome sequencing data of CD14^+^ cells from 35 RA patients is deposited in NCBI GEO with accession GSE282517. Single-cell transcriptomic data from synovial tissue of RA patients are publicly available on Synapse (https://doi.org/10.7303/syn52297840).

### 2.10. Statistical analysis

Since different methods were used for the measurement of serum IL-6, CD5L, and IGF-1 in the patient cohorts 1 and 2, the measurements were individually z-transformed to obtain comparable datasets, which were then pooled for further analysis.

Statistical analyses were performed using GraphPad Prism (version 10.1.1; GraphPad Software LLC, San Diego, CA, USA). The specific statistical tests applied are detailed in each figure legend, and relevant comparisons are shown in each graph. Differences were considered statistically significant at a P-value ≤ 0.05 (95% confidence level). Significance levels are indicated as follows: \**p* < 0.05, \*\**p* < 0.01, \*\*\**p* < 0.001, \*\*\*\**p* < 0.0001.

## 3. Results

### 3.1. CD5L delays arthritis development in CIA

CIA was induced in WT (n = 36) and CD5L^−^ (n = 39) mice by intradermal immunization with chicken CII across five independent experiments. Disease incidence, onset, and severity were monitored over 56 days. Joint swelling, the first clinical sign of arthritis, was first detected on day 23, just two days after the booster CII injection on day 21, and occurred earlier in CD5L^−^ mice than in WT controls. As shown in Table 2 and Fig. 1A, CD5L⁻ mice developed CIA more frequently than WT mice (maximum incidence 35% vs. 22%, *p* = 0.126), with the most pronounced differences during early disease (day 23, p = 0.032, and day 28, p=0.066). Arthritis onset was also earlier in CD5L⁻ mice, which exhibited a mean onset of 30.4 ± 7.5 days compared with 35.6 ± 7.5 days in WT controls (*p* = 0.08; WT: AUC = 6.958, SE = 4.962, 95% CI 0.00-16.68; CD5L⁻: AUC = 14.48, SE = 10.13, 95% CI 0.00-34.34).

**Fig. 1:**
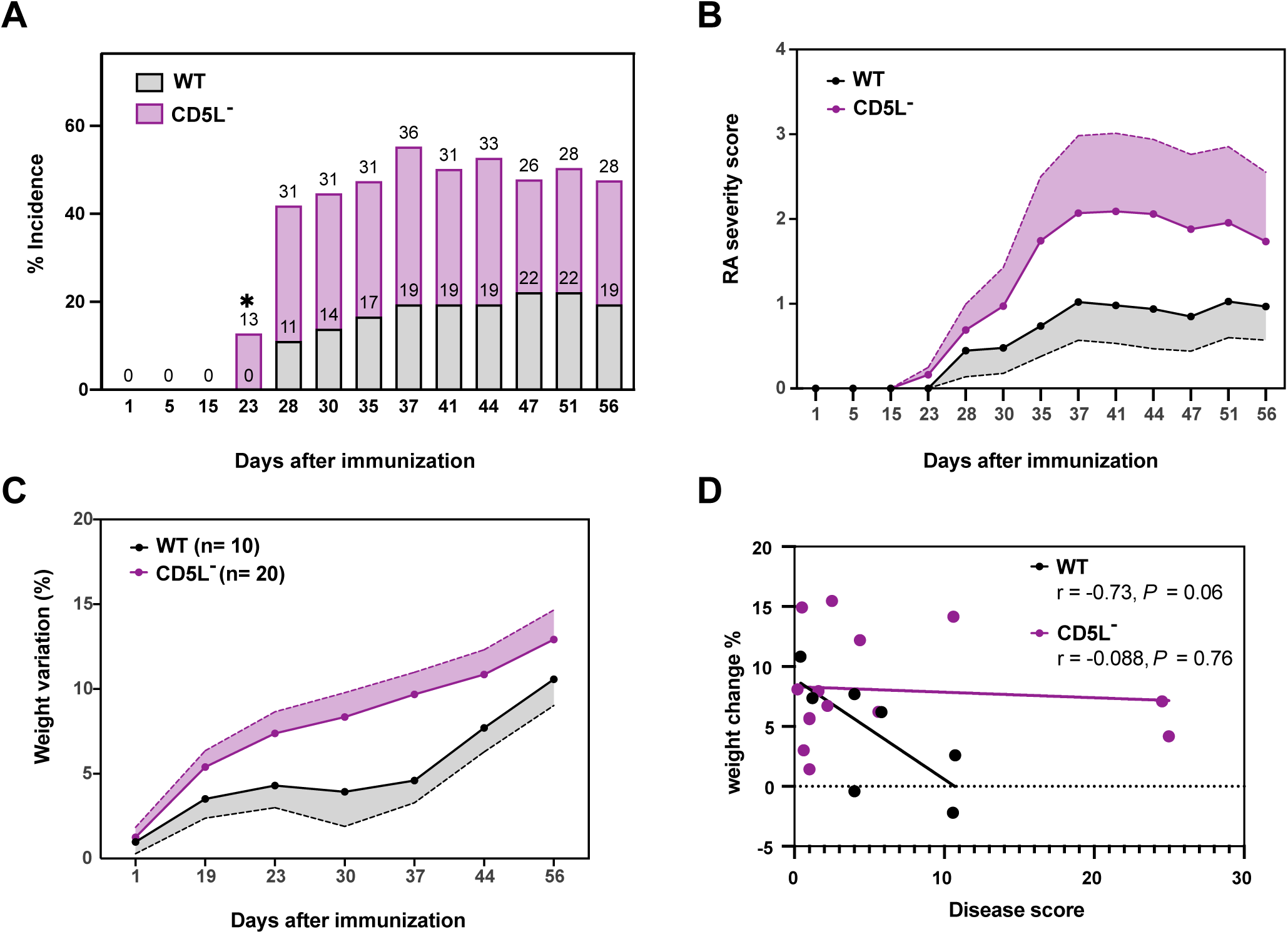
Incidence and severity of collagen-induced arthritis in WT and CD5L^−^ mice. Mice were immunized with chicken collagen on day 0 followed by a boost administration of the antigen on day 21. **A**, Incidence rates of WT and CD5L^−^ animals were compared using the Chi^2^-statistic as detailed in table 3, \**p* < 0.05. **B**, Macroscopic assessment of arthritis was performed by scoring the swelling of each paw as follows: 0 = normal, 1 = mild swelling and/or erythema, 2 = pronounced swelling, 3 = deformity and 4 = ankylosis. RA severity score of each mouse was obtained by adding the score of the four paws. **C**, Weight variation after arthritis induction. Data represent weight variation only in mice that developed arthritis, shown as mean ± SEM (n = 10 WT; n = 20 CD5L⁻), pooled from five independent experiments. B, Pearson correlation between weight variation and disease score from WT (open circles) and CD5L⁻mice (filled circles).

**Table 2:** Incidence of CIA in WT and CD5L− mice.

| Days post immunization |  | Cases | % Incidence | 95 % CI* | p<sup>#</sup> |
| --- | --- | --- | --- | --- | --- |
| 1 | WT | 0/36 | 0 | --- | --- |
|  | CD5L<sup>-</sup> | 0/39 | 0 |  |  |
| 5 | WT | 0/36 | 0 | --- | --- |
|  | CD5L<sup>-</sup> | 0/39 | 0 |  |  |
| 15 | WT | 0/36 | 0 | --- | --- |
|  | CD5L<sup>-</sup> | 0/39 | 0 |  |  |
| 23 | WT | 0/36 | 0 | 1.12-24.5 | 0.032 |
|  | CD5L<sup>-</sup> | 5/39 | 12.8 |  |  |
| 28 | WT | 4/36 | 11.1 | 1.26-40.6 | 0.066 |
|  | CD5L<sup>-</sup> | 12/39 | 30.8 |  |  |
| 30 | WT | 5/36 | 13.9 | 4.69-38.5 | 0.125 |
|  | CD5L<sup>-</sup> | 12/39 | 30.8 |  |  |
| 35 | WT | 6/36 | 16.7 | 8.09-36.3 | 0.213 |
|  | CD5L<sup>-</sup> | 12/39 | 30.8 |  |  |
| 37 | WT | 7/36 | 19.4 | 7.52-40.4 | 0.179 |
|  | CD5L<sup>-</sup> | 14/39 | 35.9 |  |  |
| 41 | WT | 7/36 | 19.4 | 11.5-34.1 | 0.330 |
|  | CD5L<sup>-</sup> | 12/39 | 30.8 |  |  |
| 44 | WT | 7/36 | 19.4 | 9.50-37.3 | 0.245 |
|  | CD5L<sup>-</sup> | 13/39 | 33.3 |  |  |
| 47 | WT | 8/36 | 22.2 | 18.8-25.6 | 0.763 |
|  | CD5L<sup>-</sup> | 10/39 | 25.6 |  |  |
| 51 | WT | 8/36 | 22.2 | 16.8-28.8 | 0.607 |
|  | CD5L<sup>-</sup> | 11/39 | 28.2 |  |  |
| 56 | WT | 7/36 | 19.4 | 13.4-31.0 | 0.440 |
|  | CD5L<sup>-</sup> | 11/39 | 28.2 |  |  |

Throughout the course of CIA, CD5L⁻ mice tended to develop more severe clinical disease, as indicated by both the number of swollen joints and the magnitude of swelling, although these differences did not reach statistical significance (Fig. 1B). Weight loss, a robust indicator of systemic inflammation in CIA, differed between WT and CD5L⁻ mice that developed arthritis (Fig. 1C), with Fig. S2A showing the day-0 weights and subsequent weight changes for the full cohort of CIA mice. On day 37, at the peak of CIA severity, weight loss strongly correlated with arthritis scores in WT mice, whereas this association was absent in CD5L⁻ mice (Fig. 1D). Histological examination of affected joints revealed dense inflammatory infiltrates, evidence of joint effusion, and, in some cases, focal loss of bone surface integrity (Fig. S2B).

### 3.2. Early CD5L changes and inflammatory cell dynamics and during CIA

To characterize the role of CD5L in CIA, we first measured serum CD5L levels in a WT cohort (n = 16, 1 developing arthritis). CD5L concentrations increased from approximately 0.81 μg/ml at day 0 to a mean of 2.65 μg/ml by day 14 (*p* = 0.0040) (Fig. 2A), then declined progressively and did not rise after the booster immunization. A modest increase was observed at day 56, without altering the overall pattern. No clinical arthritis was detected during the early CD5L surge, and when levels returned to baseline, the frequency of arthritis in WT animals resembled that of CD5L⁻ mice, which maintained undetectable CD5L throughout.

**Fig. 2:**
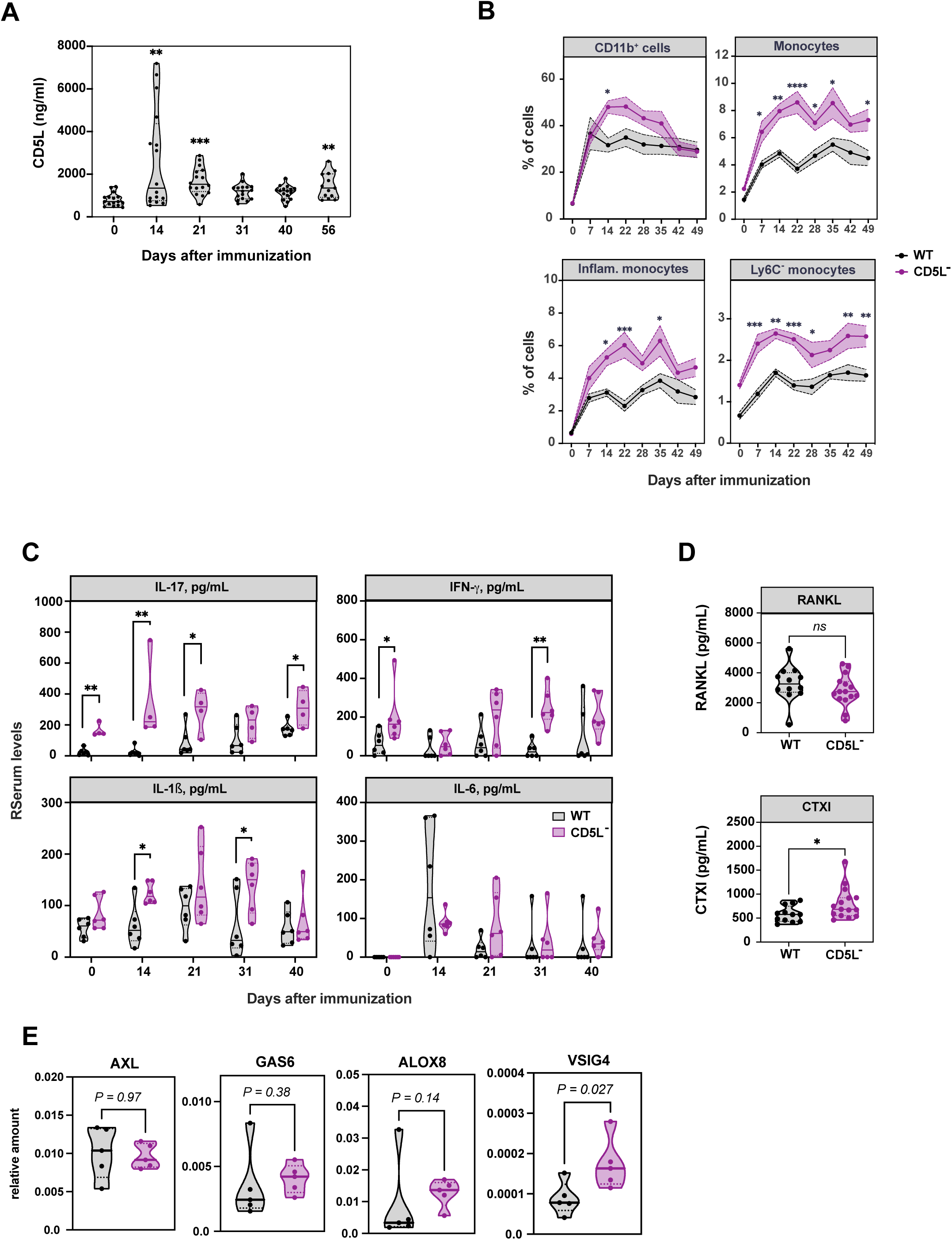
Immune profile of WT and CD5L^−^ mice upon arthritis induction. Mice were immunized with chicken collagen on day 0 followed by a boost administration of the antigen on day 21. **A**, Quantification of CD5L by ELISA in the serum of mice after immunization in the indicated time points. Statistical comparisons were made using Kruskal-Wallis test with Dunn’s multiple comparisons. **B,** The frequency of the indicated populations was analyzed in different time points after immunization in the blood of WT and CD5L^−^ mice by flow cytometry (frequencies within total live cells). Markers used for phenotyping: monocytes (Siglec-F^−^Ly6G^−^F4/80^+^CD11b^+^), inflammatory monocytes (Siglec-F^−^Ly6G^−^F4/80^+^CD11b^+^Ly6C^+^) and Ly6C^−^monocytes (Siglec-F^−^Ly6G^−^F4/80^+^CD11b^+^Ly6C^−^). Data shown are mean with SEM of n = 10 (WT) and n= 10 (CD5L^−^) mice. Statistical comparisons were made using Šídák’s multiple comparisons test. **C**, The concentration of the indicated cytokines was analyzed in different time points after immunization in the sera of WT and CD5L^−^ mice by ELISA. Statistical differences between groups were analyzed by Mann-Whitney test. The concentration of RANKL and cross-linked C-telopeptide of type I collagen (CTX-I) (**D**) was analyzed 56 days after immunization in the sera of WT and CD5L^−^ mice by ELISA. **E**, RT-qPCR quantification of the indicated genes expression, normalized with Actb (β-actin), in total spleen cells from WT or CD5L^−^ naïve mice. Mice per group: 5. Statistical differences between groups analyzed by two-tailed unpaired t-test with Welch’s correction. Data was pooled from 2 independent experiments except in c) where one representative experiment from two is depicted. Graphical representation of median and 25^th^ – 75^th^ quartiles in **A**), **C**), **D**), **E**). \**p* < 0.05, \*\**p* < 0.01, ns: not significant.

In parallel, we compared WT and CD5L⁻ mice to evaluate immune cell dynamics and systemic inflammation (n = 6, with 1 WT and 3 CD5L⁻ mice developing arthritis). CD5L⁻ mice displayed increased frequencies of circulating monocytes (Siglec-F⁻Ly6G⁻F4/80⁺CD11b⁺), affecting both classical inflammatory (Ly6C⁺) and non-classical patrolling (Ly6C⁻) subsets during the first week after CII immunization, and these elevations persisted over time (Fig. 2B). In contrast, neutrophils (Fig. S3A), as well as total CD11b⁺ myeloid cells (Fig. 2B), exhibited an early rise in CD5L⁻ mice, but these differences diminished after day 14, returning toward WT levels. Thus, while the elevation in total CD11b⁺ cells was transient, the increase in monocytes was sustained throughout the disease course. Populations of T and NK cells in the peripheral blood remained unaltered during CIA, while B cells and eosinophils were transiently decreased on day 14, with no differences between CD5L^−^ and WT mice (Fig. S3A).

Serum cytokine analyses revealed that CD5L⁻ mice had a pre-existing inflammatory background, with elevated IL-17A and IFN-γ on day 0 (Fig. 2C). Following CII immunization, IL-17A levels continued to rise, accompanied by IL-1β and IL-6. IL-1β levels correlated with the expansion of inflammatory monocytes, suggesting a potential mechanistic link. In contrast, WT mice did not exhibit a comparable cytokine surge; in the WT-only cohort analyzed earlier, the marked rise in CD5L at day 14 likely contributes to the absence of early pro-inflammatory cytokine release. Anti-CII antibody levels increased sharply after the booster at day 31 and continued rising throughout CIA, with no significant differences between genotypes (Fig. S3B).

To assess the impact of CD5L deficiency on bone remodeling, we measured serum markers of bone resorption. CD5L⁻ mice showed elevated CTX-I levels, whereas RANKL remained similar to WT controls (Fig. 2D). To explore mechanisms linking inflammation and osteoclast activity, we analyzed the expression of *GAS6*, *AXL*, *ALOX8*, and *VSIG4*, genes involved in anti-inflammatory signaling and the regulation of monocyte/macrophage activity. In CD5L⁻ mice, *AXL* expression was reduced, while *GAS6* and *ALOX8* were modestly increased, and *VSIG4* showed the strongest elevation, indicating a shift in regulatory pathways that may influence osteoclast differentiation and limit excessive bone loss (Fig. 2E).

Overall, the early rise in serum CD5L in WT mice appeared to confer protection during CIA, whereas CD5L⁻ mice showed elevated baseline IL-17 and IFN-γ, leading to sustained inflammatory monocyte expansion and more severe arthritis. The gene expression profile in CD5L⁻ mice further reflected this heightened inflammatory state, with reduced *AXL* but compensatory increases in *GAS6*, *ALOX8*, and especially *VSIG4*, indicating attempts to counterbalance persistent immune activation.

### 3.3. CD5L promotes an anti-inflammatory phenotype in human monocytes

To provide experimental evidence linking CD5L to the regulation of monocyte phenotypes, PBMCs (n = 8) were activated with LPS and ConA and subsequently exposed to recombinant CD5L (rCD5L; 1 μg/ml), or CD5L + IgG, for 24 h before flow cytometric analysis. Gating on CD4⁻CD11b⁺ monocytes, we found that CD5L treatment reduced surface expression of the αM-integrin CD11b, as measured by MFI (*p* = 0.031). Furthermore, the combined CD5L + IgG condition decreased the expression of integrin β1 (CD29) on CD14⁺ monocytes (*p* = 0.031), indicating an enhanced regulatory effect that may further limit monocyte migratory and tissue-infiltrative capacity (Fig. 3A).

**Fig. 3:**
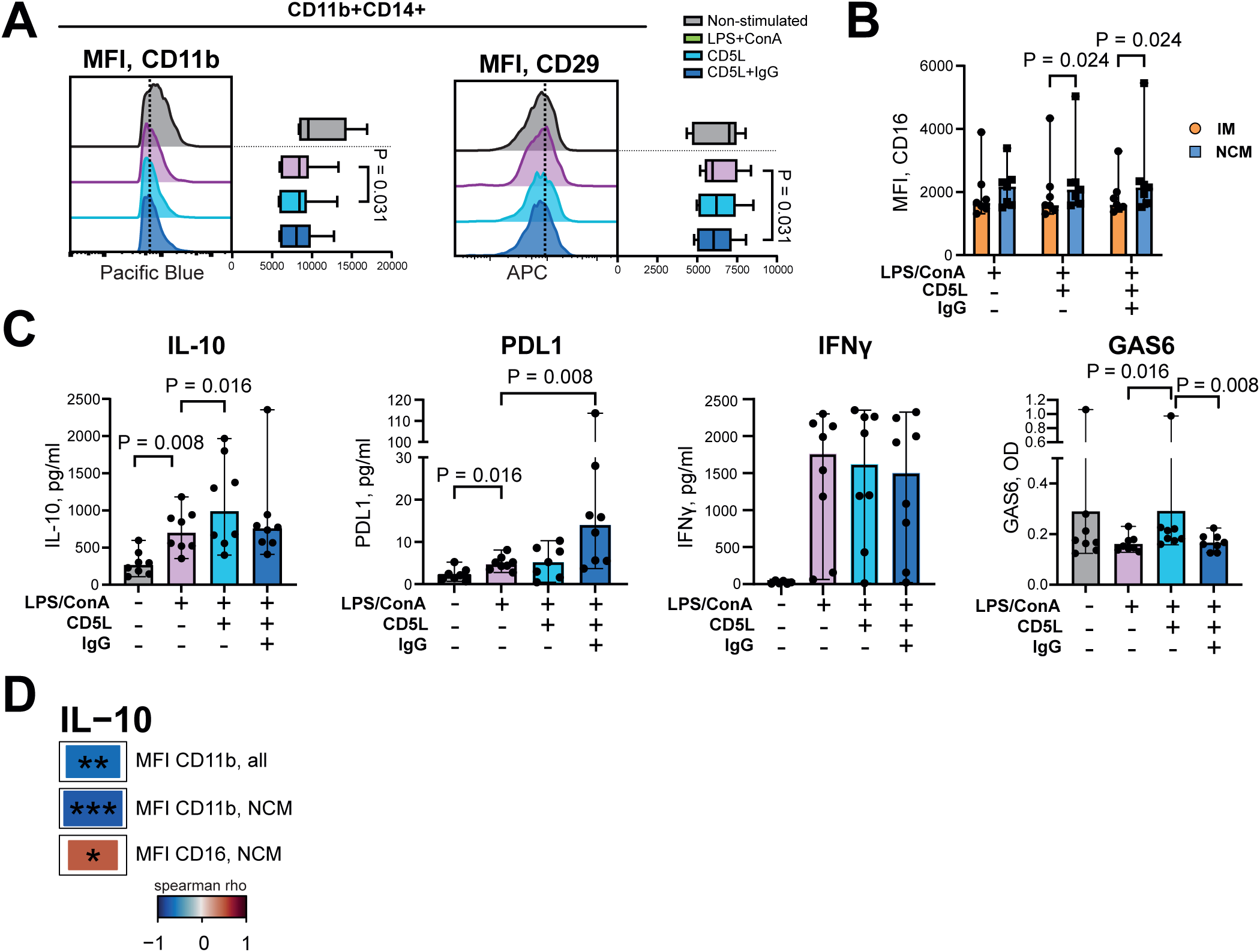
CD5L induces an anti-inflammatory phenotype in human monocytes. **A**, Representative histograms showing Pacific Blue-CD11b and APC-CD29 expression in human CD11b⁺CD14⁺CD4⁻ peripheral blood leukocytes stimulated with LPS and ConA for 48 h, with the addition of human recombinant CD5L (1 μg/ml) or recombinant CD5L plus human IgG (equimolar) during the final 24 h. Expression was assessed by flow cytometry. Box plots depict mean fluorescence intensity (MFI) of CD11b and CD29 in cultured cells (n = 8). P values were calculated using a paired Wilcoxon test. **B**, Column plots of CD16 expression on intermediate (IM, CD14^+^CD16^+^) and non-classical (NCM, CD14^low^CD16^+^) monocyte subsets. **C**, Protein levels of IL-10, PDL1, IFN-γ and GAS6 in the supernatants of cell cultures. **D**, Heat maps showing Spearman correlation (ρ) values between IL-10 concentrations in culture supernatants and the expression intensity of CD11b and CD16, as measured by flow cytometry. Statistical significance is indicated as \**p* < 0.05; \*\****p*** < 0.01; \*\*\****p*** < 0.001.

Analysis of the major monocyte subsets – classical, intermediate, and non-classical (gating strategy shown in Fig. S4A) – revealed no significant effect of CD5L stimulation (Fig. S4B). However, among the intermediate and non-classical subsets, CD5L increased the expression intensity of CD16/FCGRIIIA, shifting the balance toward the non-classical monocyte subset (Fig. 3B). Surface expression of the TLR4 receptor CD14 was unaffected by CD5L stimulation.

Measuring cytokine production in supernatants of CD5L-stimulated cells, we found a significant increase in protein production of IL-10 (*p* = 0.016), GAS6 (*p* = 0.016), and PDL1 (CD5L + IgG, *p* = 0.008), but no significant effect on the production of IFN-γ as compared to LPS + ConA alone (Fig. 3C). Correlating IL-10 production to monocyte phenotypes, we found a negative correlation with CD11b expression on CD4^−^CD11b^+^ monocytes (Spearman ρ = –0.616, *p* = 0.003). On non-classical CD14^low^CD16^+^ monocytes, the correlation between IL-10 and CD11b was stronger (ρ = –0.729, *p* = 0.0002); additionally, a positive correlation to CD16 expression (ρ = 0.511, *p* = 0.018) was observed (Fig. 3D).

Taken together, these results indicate that CD5L exerts anti-inflammatory activity by suppressing integrin expression and promoting IL-10 and PD-L1 production, a profile associated with increased proportions of non-classical monocytes.

### 3.4. Histone acetylation-dependent production of CD5L by human PBMCs

To examine CD5L production by human monocytes, we stimulated freshly prepared PBMC cultures (n = 20) with LPS + ConA, and subsequently stimulated them with Fcγ-receptor ligand IgG, class I HDAC inhibitor (HDACi, valproate), proteasome inhibitor (PI, bortezomib) and Lyn agonist (LynA, talimidone), to induce an anti-inflammatory response. We found that CD5L protein levels were not increased significantly after 48 h of stimulation with LPS + ConA, while addition of HDACi to LPS + ConA-stimulated PBMCs demonstrated a significant rise in CD5L production (*p* = 0.0001) (Fig. 4A). CD5L production remained unchanged/low after cell stimulation using human IgG, LynA and PI (Fig. S4C). The HDACi-induced increase of CD5L production coincided with a significant increase of IL-10 production (*p* = 0.011) and a decrease of IFN-γ (*p* = 0.002), GAS6 (*p* = 0.067), and PD-L1 (*p* = 0.008) (Fig. 4A). LynA affected neither CD5L nor IFN-γ production by PBMC cultures stimulated with HDACi (Fig. S4D). Correlation analyses showed that IFN-γ production had an inverse association with CD5L (ρ = –0.494, *p* = 0.004), IL-10 (ρ = –0.541, *p* = 0.0014) and CD16 expression (ρ = –0.469, *p* = 0.010. Fig. 4B).

**Fig. 4:**
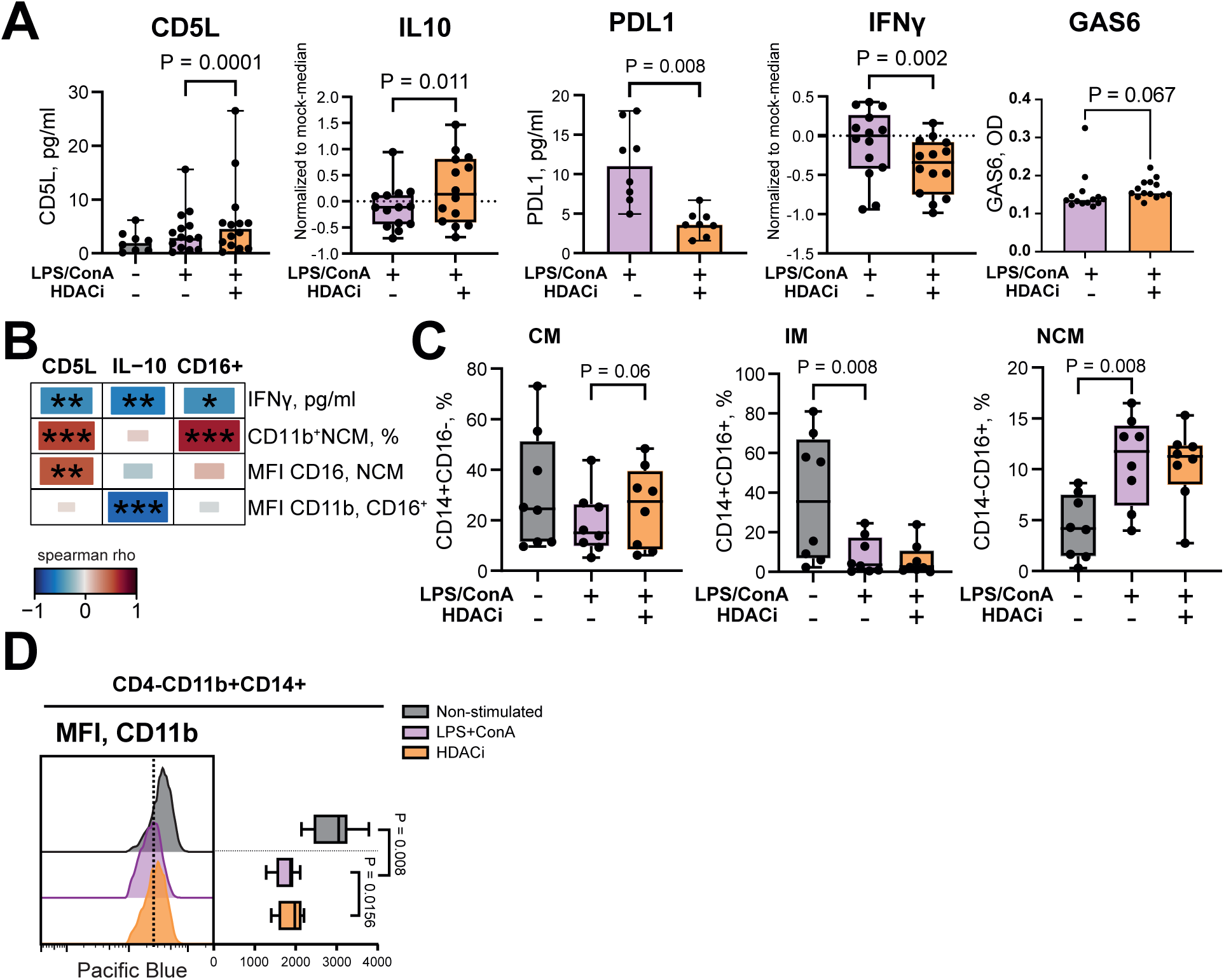
HDAC inhibition promotes CD5L production and skews human monocytes toward an anti-inflammatory phenotype. **A**, Box plots showing CD5L, IL-10, PD-L1, IFN-γ, and GAS6 protein levels in supernatants from human peripheral blood leukocyte cultures (n = 14) stimulated with LPS and ConA (5 μg/mL and 0.625 μg/m, respectively) for 48 h, with the addition of a histone deacetylase inhibitor (HDACi, 50 μg/ml) during the final 24 h. **B**, Heat maps displaying Spearman correlation (ρ) values between CD5L and IL-10 levels in supernatants, CD16 expression intensity measured by flow cytometry, IFN-γ production, the relative size of the non-classical monocyte (NCM) subset, and the mean fluorescence intensity (MFI) of CD16 and CD11b. Statistical significance is indicated as \**p* < 0.05; \*\****p*** < 0.01; \*\*\****p*** < 0.001. **C,** Box plots illustrating changes in classical (CM; CD14⁺CD16⁻), intermediate (IM; CD14⁺CD16⁺), and non-classical (NCM; CD14^low^CD16⁺) monocyte subsets in stimulated human leukocyte cultures (n = 14), treated as described in panel (**A**). **D,** Representative histograms of Pacific Blue-CD11b expression in human CD11b⁺CD14⁺CD4⁻ blood leukocytes stimulated as indicated and analyzed by flow cytometry. Box plots summarize CD11b mean fluorescence intensity (MFI) across conditions. P values were calculated using a paired Wilcoxon test.

To evaluate how the HDACi-induced CD5L production was related to the change of monocyte phenotype, we used flow cytometry. HDACi stimulation favored the increase of classical (CD14^+^CD16^−^) monocytes (*p* = 0.0625), having no significant effect on the intermediate and non-classical subsets (Fig. 4C). HDACi increased α_M_-integrin CD11b expression within the CD4^−^CD11b^+^ gate (*p* = 0.016) (Fig. 4D).

Correlation analyses showed that CD5L levels in HDACi-treated cells correlated positively to both the population size of non-classical CD14^low^CD16^+^ monocytes (Spearman ρ = 0.582, *p* = 0.0009) and CD16 expression intensity on this subset (ρ = 0.529, *p* = 0.003), which collectively suggested a connection between non-classical monocyte activation and CD5L production (Fig. 4B). IL-10 levels in HDACi-treated cell cultures were negatively correlated to CD11b MFI on CD16^+^ monocytes (Fig. 4B), recapitulating the pattern previously observed following CD5L stimulation (Fig. 3D). Since CD5L stimulation increased IL-10 and GAS6 production, these mediators may act as intermediaries through which CD5L exerts its effects on PBMCs.

Taken together, these findings support an HDAC-dependent mechanism for CD5L production, acting in conjunction with IL-10 and GAS6 to counter the accrual of the pro-inflammatory monocyte subset.

### 3.5. CD5L production delineates the efferocytosis phenotype of CD14^+^ cells

To further characterize the CD5L-producing CD14^+^ cells, we measured CD5L levels in supernatants of LPS-activated CD14^+^ cell cultures of 35 RA patients (Table 1). CD5L protein concentrations exceeding twice the assay detection limit (0.6 pg/ml) were measurable in the supernatants of 10 out of 35 CD14⁺ cell cultures, hereafter designated as CD5L^hi^CD14⁺ cells. These cells displayed significantly higher IL-10 protein levels (27.5 vs. 22 pg/ml, *p* = 0.0255) and *IL-10* mRNA expression (2262 vs. 133 relative units, *p* = 0.0028) (Fig. 5A, 5B). Furthermore, RA patients with CD5L^hi^CD14^+^ cells exhibited significantly higher PD-L1 levels (25.5 vs. 18.87 pg/ml, *p* = 0.0304), elevated white blood cell counts (6.85 vs. 5.2 × 10⁹/l, *p* = 0.051) and platelet counts (293 vs. 225 × 10⁹/l, *p* = 0.0031), as well as increased serum IFN-γ levels (19.60 vs. 17.02 pg/ml, *p* = 0.064) (Fig. 5A). No differences were observed in the levels of IFN-γ, IL-1β, IL-8, and IL-6 between the supernatants of CD5L^hi^CD14^+^ and CD5^low^CD14^+^ cells (Fig. S5A).

**Fig. 5:**
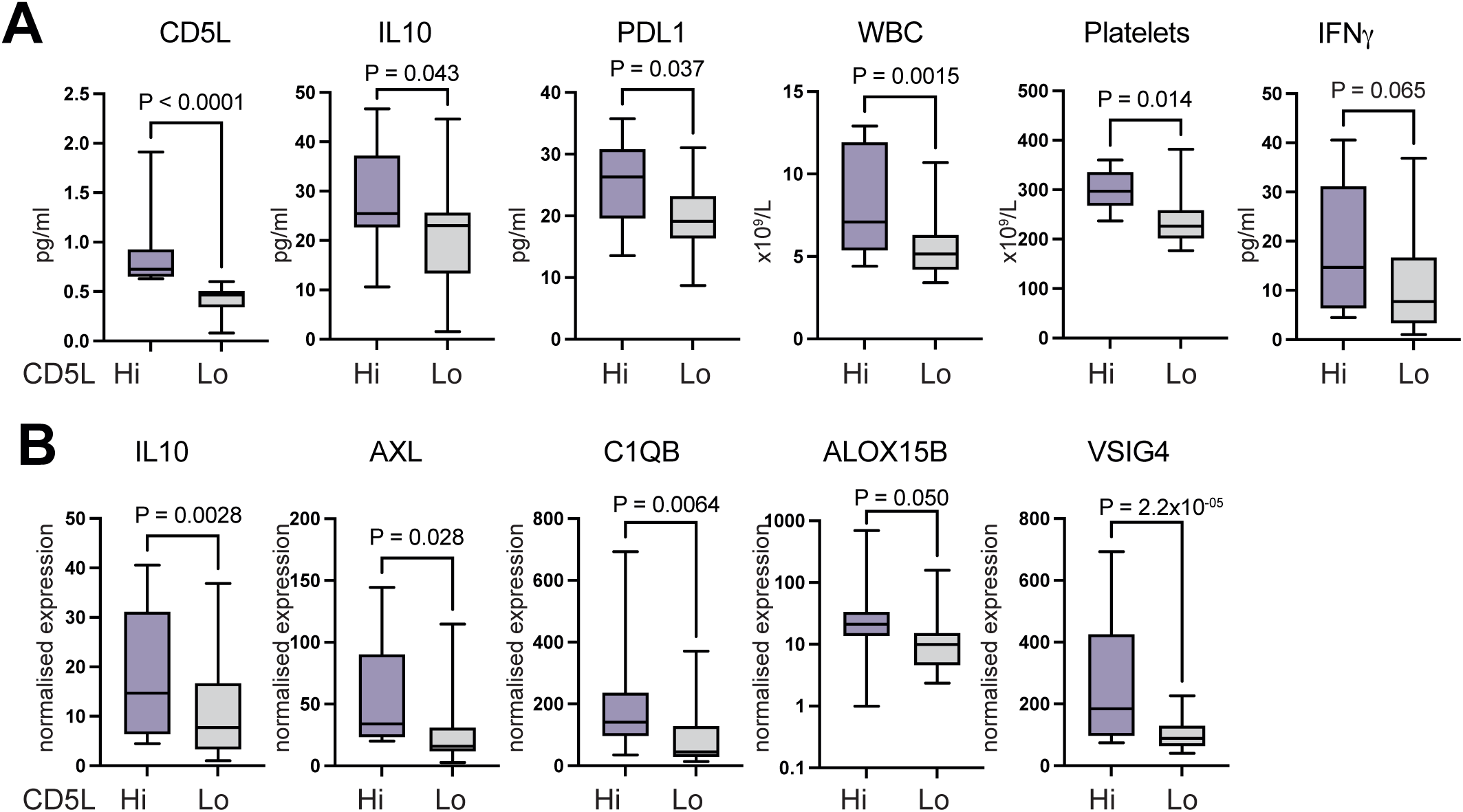
CD5L production identifies efferocytosis-prone CD14⁺ monocytes in RA patients. CD14⁺ cells were isolated from peripheral blood leukocyte cultures of 35 RA patients, and CD5L levels in culture supernatants were measured by ELISA. Cultures with CD5L protein levels above 0.6 pg/mL (double detection limit) were classified as CD5L^hi^, and the remaining as CD5L^low^. **A,** Box plots showing protein levels of CD5L, IL-10, and PD-L1 in supernatants, as well as patient white blood cell (WBC) and platelet counts, and serum IFN-γ levels, comparing CD5L^hi^ (n = 10) and CD5L^low^ (n = 25) CD14⁺ cells. **B,** Box plots of normalized gene expression levels in CD14⁺ cells measured by RNA-seq. P values were calculated using DESeq2; nominal p values are indicated.

The high IL-10 production in CD5L^hi^CD14^+^ cells was accompanied by transcriptional upregulation of the C1q subunits *C1QA*, *C1QB*, and *C1QC*, which opsonize apoptotic cells and promote efferocytosis [53], as well as the TAM receptor *AXL* and pro-resolving regulators *ALOX15B* and *VSIG4*, suggesting that CD5L-producing monocytes acquire an efferocytosis-prone phenotype (Fig. 5B). In agreement with this, the pathway enrichment analysis of the genes functionally dependent on high CD5L production in CD14^+^ cells identified negative regulation of cytokine production and control of immune system processes and cell motility as the top biological processes associated with high CD5L production in CD14^+^ cells (Fig. S5B). Among the suppressed biological processes in CD5L^hi^CD14^+^ cells, the downregulation of ATPase– and GTPase-dependent hydrolase activity was the most prominently enriched process, showing an inverse relationship with CD5L production in CD14⁺ cells.

Taken together, transcriptome analysis of CD5L^hi^CD14^+^ cells revealed the acquisition of an efferocytosis-prone phenotype and enhanced IL-10 production. These findings mirrored the results observed in HDACi-treated cultures, suggesting that CD5L production by monocytes is associated with suppression of their pro-inflammatory effector functions.

### 3.6. Serum CD5L levels shape transcriptome of blood CD14^+^ cells

To further explore the effects of extracellular CD5L on RA monocytes in vivo, we analyzed the transcriptome of CD14⁺ cells in relation to serum CD5L levels. Monocyte subset composition was estimated in silico using the normalized sum of RNA-seq expression for subset-specific marker genes identified in a recent single-cell transcriptome study [54]. IL-1β–rich monocytes exhibited the highest cumulative marker gene expression, followed by classical and IFN-γ-imprinted monocytes, while non-classical monocytes showed the lowest expression levels (Fig. 6A, 6B). Among the genes upregulated in proportion with increasing serum CD5L levels, we identified the genes of non-classical monocytes including *FCGR3A, MT2A, C1QA, C1QB*, and *LILRA5* (Fig. 6C), which reproduced the experimental findings of CD5L-dependent enrichment for CD16^+^ monocytes. Additionally, the signature genes of the IFN-imprinted monocytes, including *ISG15*, *TYMP*, *FTH1*, and also *IFNG*, *JUN*, *CEBPB* and *CBX6* genes, were upregulated in parallel with increasing serum levels of CD5L (Fig. 6C).

**Fig. 6:**
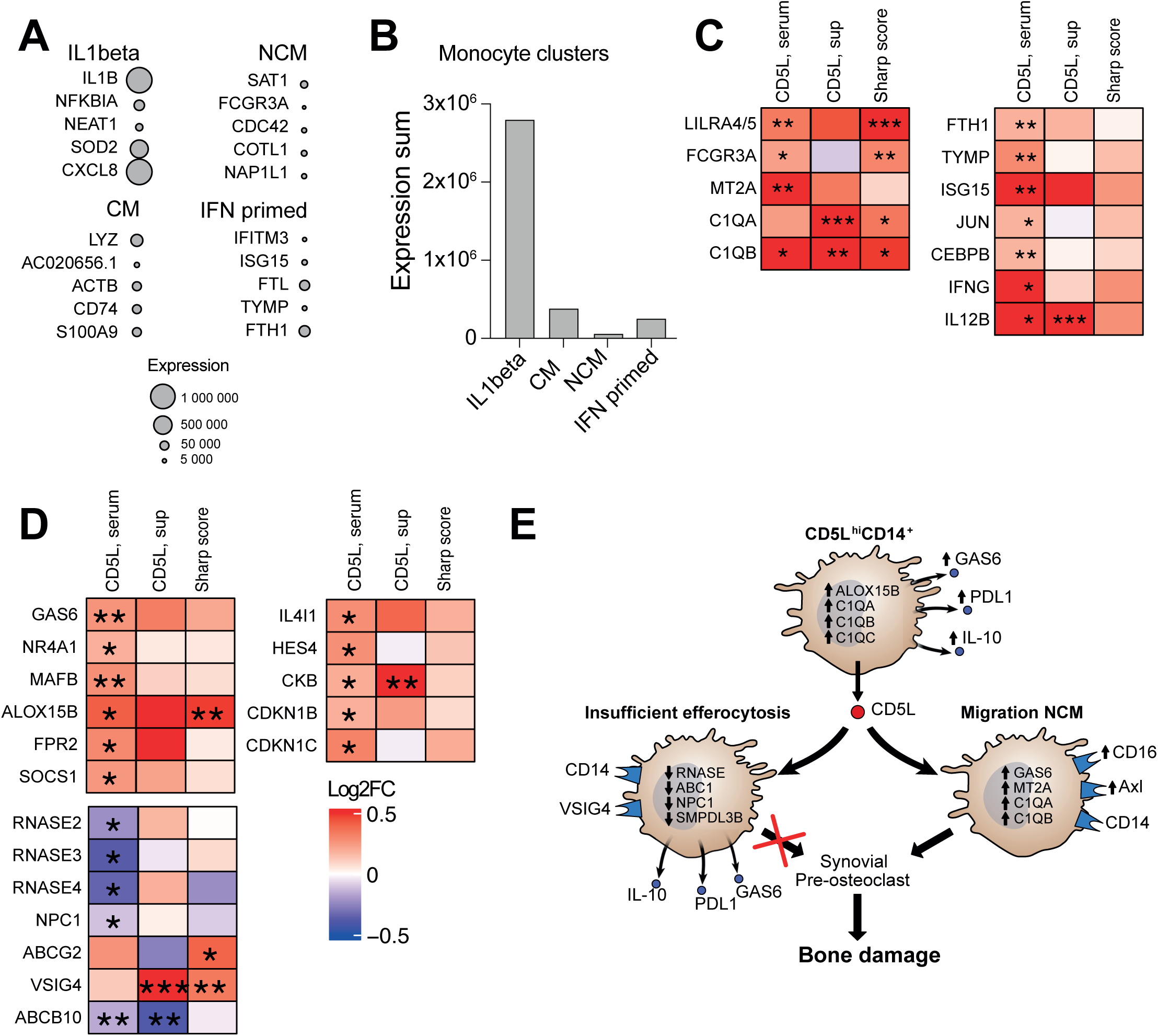
CD5L shapes the transcriptional profile of CD14⁺ monocytes in RA patients. CD14⁺ cells were isolated from peripheral blood leukocyte cultures of 35 RA patients, activated with LPS for 2 h, and subjected to RNA sequencing (RNA-seq, Illumina). **A,** Bubble plot showing mean normalized expression of gene markers for four monocyte clusters identified in human blood leukocytes. **B,** Bar plot representing the distribution of monocyte clusters based on the sum of cluster-specific gene expression. **C,** Heat map of gene expression changes (log₂ fold change) in non-classical and IFN-primed monocyte clusters in regression to serum CD5L levels and joint skeletal damage quantified by vdH-Sharp score. P values were calculated using DESeq2; nominal P values are indicated as \**p* < 0.05; \*\****p*** < 0.01; \*\*\****p*** < 0.001. **D,** Heat map of gene expression changes (log₂ fold change, FC) for efferocytosis markers (1), resolution mediators (2), and immunoregulatory macrophage program genes (3) in regression to serum CD5L levels and vdH-Sharp score. **E,** Schematic representation of a monocyte conditioned by CD5L.

Further analysis of the CD14⁺ cell transcriptome in relation to serum CD5L levels revealed the accumulation of genes consistent with the pro-efferocytosis phenotype observed in CD5L^hi^CD14^+^ cells. This included increased expression of *C1QB* and *CXCL16*, which opsonize apoptotic cells and PS-coated bodies; transcription factors *MAFB* and *NR4A1*, which govern anti-inflammatory molecules *NFKBID* and *NR4A3* that repress the NF-κB pathway; upregulation of the TAM receptor ligand *GAS6*, mediating *SOCS1* production; and regulators of inflammation resolution such as *ALOX15B* and *FPR2*, potentially enhancing efferocytosis capacity (Fig. 6D).

However, several lysosomal proteins involved in apoptotic cargo processing were downregulated with increasing serum CD5L levels, including the RNASE genes, multiple ABC1 lipid exporters, the cholesterol exporter *NPC1*, and the internalization-related *SMPDL3B*, suggesting insufficient or overloaded efferocytosis (Fig. 6D). Supporting this, immunoregulatory macrophage program genes, including *HES4*, *IL4I1*, *CKB*, *CDKN1B/CDKN1C*, and *VSIG4* genes, previously identified in tumor-conditioned macrophages [55, 56], were upregulated, indicating dampened resolution biochemistry and impaired apoptotic cell clearance (Fig. 6D).

Taken together, these findings indicate that serum CD5L levels shape the transcriptome of CD14⁺ cells in RA, promoting the formation of efferocytosis-prone monocytes observed in experimental studies, while also contributing to the accumulation of non-classical, IFN-primed monocytes with potentially harmful intra-articular effects, as summarized schematically in a CD5L-conditioned monocyte model (Fig. 6E).

### 3.7. Serum CD5L levels are associated with increased joint skeletal damage in RA

Serum CD5L levels were measured in 80 RA patients and 11 healthy volunteers (Table 1). We found that serum CD5L levels tended to be higher in RA patients than in controls, although our results did not reach statistical significance (*p* = 0.057; Fig. 7A, left). Comparing paired samples of serum and synovial fluid of 5 RA patients, we found a significant accumulation of CD5L levels in the joint cavity of RA patients (*p* < 0.05; Fig. 7A, right), consistent with a previous report [57].

**Fig. 7:**
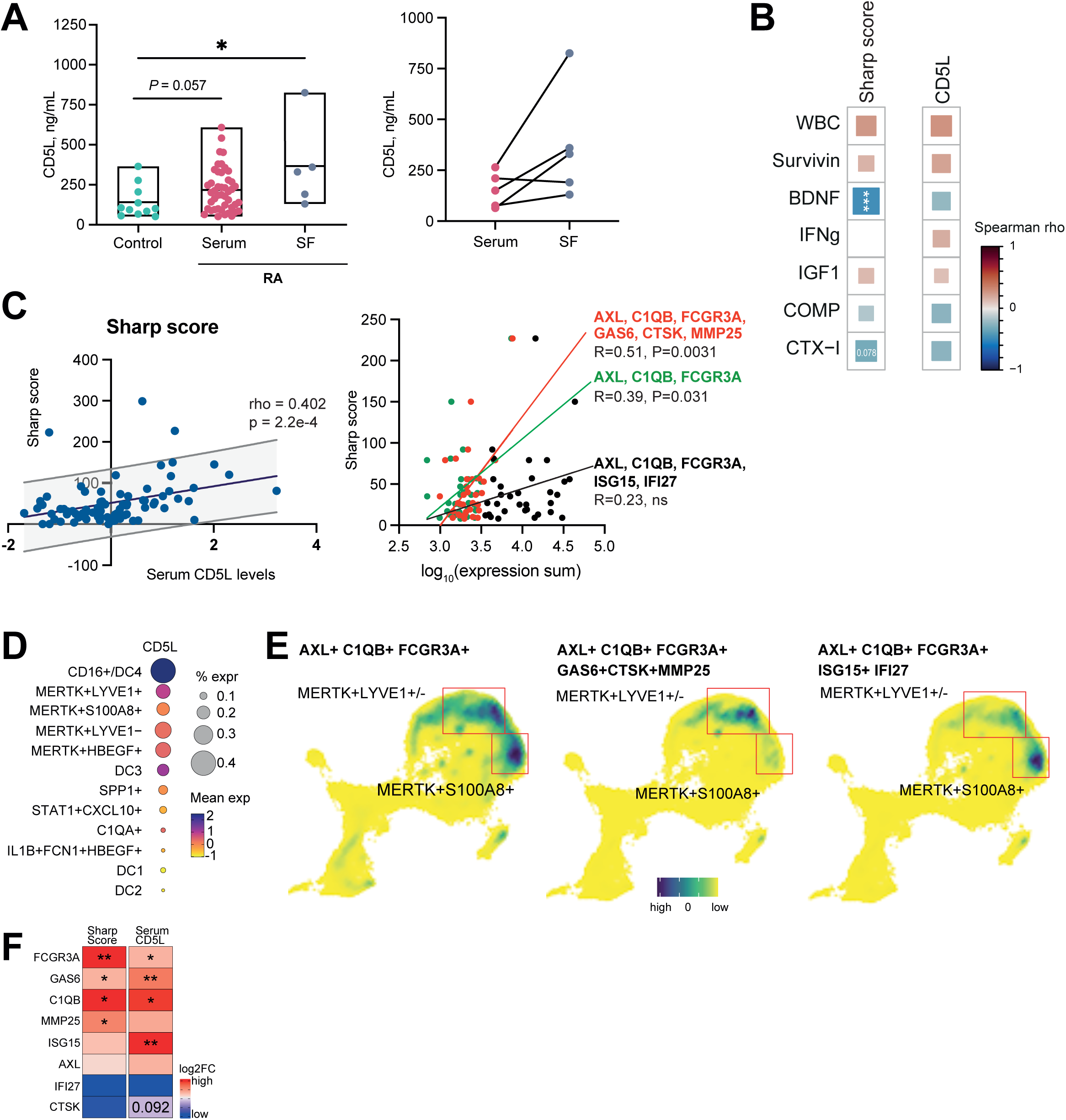
Low CD5L expression in synovial macrophages associates with joint damage in RA. **A**, Box plots of CD5L protein levels in serum of RA patients and healthy controls (1), paired serum and synovial fluid samples from RA patients (2), and anti-CD5L antibodies in serum of RA patients and controls (3). **B,** Heat map of Spearman correlations (ρ) between serum CD5L levels or vdH-Sharp joint damage scores and serum cytokine and growth factor levels. Statistical significance is indicated as \**p* < 0.05; \*\****p*** < 0.01; \*\*\****p*** < 0.001. **C,** Dot plots showing correlations between serum CD5L levels and radiographic joint damage in 80 RA patients (1), and correlation of a CD5L-dependent gene signature in non-classical monocytes with vdH-Sharp score (2). Enrichment with osteoclast-related markers enhanced the correlation, whereas inclusion of IFN-sensitive genes abolished it. **D,** Bubble plot of the percentage frequency of CD5L⁺ cells within synovial myeloid clusters. **E,** UMAP of the scaled expression intensity sum of the CD5L-dependent signature in synovial myeloid cells from single-cell transcriptomics. Further enrichment distinguishes IFN-primed versus osteoclast-fostering macrophages. **F,** Heat map of a CD5L-dependent osteoclastogenic signature in blood CD14⁺ cells, showing a positive correlation with vdH-Sharp joint damage scores.

We then assessed the correlation between CD5L serum levels and clinical data of RA patients. This analysis revealed a positive correlation between CD5L levels and RA severity, assessed by clinical disease activity (DAS28, ρ = 0.31, *p* = 0.006), and joint skeletal damage (JSD) quantified by vdH-Sharp score (ρ = 0.402, *p* = 0.00022) (Fig. 7B, C). There was also an inverse correlation between CD5L and serum COMP (ρ = –0.30, *p* = 0.026) and CTX-I levels (ρ = –0.32, *p* = 0.078), acknowledged biomarkers of bone and cartilage metabolism [58, 59].

Linear regression analysis revealed a significant association between serum CD5L levels and vdH-Sharp scores in radiographs of hands and feet (Fig. 7C). Analysis of JSD in the 35 RA patients, the source of the CD14⁺ cells used in this study, showed correlations between vdH-Sharp scores and disease duration (Spearman ρ = 0.503, *p* = 0.004), as well as with serum levels of IL-8 (ρ = 0.369, *p* = 0.041), BDNF (ρ = – 0.470, *p* = 0.008), and RANKL (ρ = –0.346, *p* = 0.057).

To explore relationship between the radiographic damage and CD5L-dependent transcriptome of CD14^+^ cells, we compared the genes transcription changed in relation to these two parameters. The pathway enrichment analysis identified the genes functional in leukocyte activation, cell adhesion, and cell motility that were significantly upregulated both in proportion to vdH-Sharp score and to serum CD5L levels (Fig. S5C). Namely, in the regression analysis, both serum CD5L levels and the vdH–Sharp score were associated with markers of efferocytosis-prone non-classical monocytes, including *C1QB*, *C1QA*, *GAS6*, *ALOX15B*, *VSIG4*, as well as *FCGR3A*, the *C1Q* subunits, and the *LILRB4/5* receptors, but not with genes characteristic of IFN-primed monocytes (Fig. 7C).

### 3.8. Scant CD5L expression in synovial macrophages facilitates joint damage in RA

To assess CD5L-dependent effects in resident monocytes of synovial tissue, we analyzed the myeloid cell clusters (n = 76,181 cells) previously characterized in RA synovia [50]. In total, non-zero *CD5L* expression was detected in only 0.17% (129 out of 76,181) of myeloid cells in RA synovia. Of these *CD5L*-expressing cells, 62% (80 out of 129) were within the MerTK⁺ clusters, specifically the MerTK⁺S100A8⁺, MerTK⁺SELENOP⁺LYVE1⁻, and MerTK⁺SELENOP⁺LYVE1⁺ populations, whereas only 0.05% (6 out of 129) corresponded to monocyte-derived CD16⁺ dendritic cells, known for their high migratory potential (Fig. 7D). This pattern suggested that either predominantly CD5L-deficient monocytes traffic into the RA synovium or, alternatively, that *CD5L* expression is downregulated after macrophages enter the synovial tissue. Consequently, the CD5L imprint on synovial macrophages is likely driven by CD5L availability in the serum or synovial fluid rather than by local production within the joint.

In macrophages and dendritic cells, the TAM family of receptor tyrosine kinases are represented by expression of AXL and MerTK. In experimental arthritis, both AXL and MerTK have anti-inflammatory properties acting in the FCGR3A rich setting [60]. In RA synovial tissue, we found that the CD5L-dependent monocyte signature consisting of *AXL-FCGR3A-C1QB* genes was enriched within the CD5L-expressing and, potentially, inflammation resolving MerTK^+^LYVE1^+^ cluster [35] and the pro-inflammatory and joint damaging MerTK^+^S100A8^+^ cluster [61] (Fig. 7E). These cells also possessed high expression of the TAM receptor ligand GAS6 which was upregulated in the blood CD14 transcriptome in direct proportion to JSD quantified by the vdH-Sharp score and upregulated in connection with increasing levels of serum CD5L. The MerTK^+^S100A8^+^ cluster was marked with IFN-sensitive genes ISG15, IFI27, which were CD5L-dependent and identified the IFN-primed non-classical monocytes. In contrast, expression of GAS6 and NRA41 recognized MerTK^+^LYVE1^+^ cluster (Fig. 7E).

To elucidate a connection between AXL, C1QB, and FCGR3A to JSD, we analyzed expression of the osteoclast progenitor markers *CTSK*, coding for cathepsin K, and MMP25 in the defined MerTK^+^LYVE1^+^ and MerTK^+^S100A8^+^ clusters. We found that CTSK^+^MMP25^+^ pro-osteoclasts were enriched within the MerTK^+^LYVE1^+^ cluster and combined with GAS6 gene expression. The expression sum of *AXL, FCGR3A, C1QB, GAS6, CTSK, MMP25* genes in blood CD14^+^ cells had a strong positive correlation to the vdH-Sharp score of RA patients (Fig. 7F). Interestingly, the CD5L-dependent markers of post-engulfing processing and serial efferocytosis ALOX15B, FPR2/3, RENASE2, VSIG4, NPC1, ABCG1/2, HES4, were not expressed in MerTK^+^ clusters of RA synovia reflecting either absence of CD5L expressing cells or low intensity of tissue efferocytosis.

Overall, this functional analysis confirmed that serum CD5L levels were positively related to the skeletal joint damage caused by RA. Scant CD5L-expressing cells in RA synovia could be a reason for the expansion of AXL^+^FCGR3A^+^GAS6^+^ macrophages, which harbored osteoclast progenitors with established joint damaging properties.

## 4. Discussion

In this study, we demonstrated that the exposure to CD5L and CD5L production in CD14^+^ cells had anti-inflammatory effect by inducing production of IL-10 and PD-L1 and lowering surface integrins, and inflammation resolving effect by promoting shift to monocytes engaged in efferocytosis both in culture and in RA patients. In RA patients, increasing serum CD5L levels were associated with the acquisition of a complete efferocytosis program, characterized by high expression of complement C1q subunits, activation of the GAS6/TAM receptor pathway, and upregulation of the resolvin regulators ALOX15B and FPR2. This profile is consistent with the efferocytic potential of monocytes exposed to, or producing, CD5L.

In parallel with inflammation resolving features, the CD5L-treated leukocyte cultures enrich for FCGR3A/CD16+ monocytes, which in RA patients was associated with accumulation of IFN-primed and non-classical monocytes. The pro-inflammatory program activated by IFN-priming is expected to reduce efferocytosis capacity [62]. Additionally, development of two other processes counteracted efficient efferocytosis in RA monocytes that were transcriptional depression of resolution/efflux machinery (ABC1 family genes, NPC1, RNASEs) and upregulation of immunoregulatory genes (HES4, IL4i1, CKB, CDKN1B/CDKN1C, VSIG4) leaving a free way for the IFN-primed non-classical monocytes into RA joints.

The bulk transcriptome of CD14^+^ cells was used in the exploratory part of the study, thus development of anergic/ low metabolic monocytes and enrichment for the tissue trafficking monocyte subset could present two parallel and even competing events ongoing in different cells. We addressed this notion by analyzing a vast single-cell data set of RA synovia. The analysis unexpectedly demonstrated a profound CD5L deficiency of synovial myeloid cells. In contrast, the TAM-expressing synovial clusters were heavily enriched with non-classical (FCGR3A^+^C1QB^+^AXL^+^) cells fostering osteoclast precursors. Moreover, a CD5L-dependent signature including expression of FCGR3A, C1QB, AXL, GAS6, MMP25 and CTSK in blood monocytes of RA patients correlated with their joint skeletal damage.

The analysis of transcriptome of synovial tissue myeloid cell clusters demonstrated a striking scarcity of CD5L positive cells despite the abundance of CD5L in synovial fluid. Those few CD5L^+^ cells were largely found within MerTK^+^ clusters of myeloid synovial cells. Despite the low number of CD5L expressing cells, the MerTK^+^ clusters had a strong representation of non-classical FCGR3A^+^C1QB^+^AXL^+^ cells, which expanded with CD5L exposure.

The joint damaging properties of non-classical CD16^+^ monocytes have been demonstrated due to their ability of osteoclast differentiation in mouse experimental arthritis models [63, 64]. We offer a further step in this direction and demonstrated that enrichment with non-classical monocyte subsets harbored features of environment dependent adaptation in response to external signals. Co-expression of GAS6 with TAM receptors signify is a characteristic feature of such joint-damaging monocytes. TAM receptors AXL and MerTK are negative regulators of bone healing. Axl inhibition prevents osteoclast differentiation [65], while GAS6 has a bone destructive role in mice periodontitis [66]. MerTK inhibition increased osteoblastogenesis. On the other hand, no signs of osteoclast inclination were found in the IFN-primed FCGR3A^+^C1QB^+^AXL^+^ synovial macrophages consistent with their high motility and adhesion profile described in systemic sclerosis [67].

CD5L increase occurred in the pre-arthritis and early arthritis phases of CIA in WT mice. No clinical arthritis was registered in WT mice during the period corresponding to CD5L increase. Consistent with previous reports of anti-inflammatory propertied of CD5L [6, 7], CD5L-deficiency in CIA mice was associated with a remarkable efflux of monocyte/macrophages into the blood [68], and excessive production of IL-1β, IL-17, and IFN-γ [11]. To simplify the role of CD5L in RA, we performed our studies on CD5L-stimulated primary cell cultures and assessed the transcriptional effects of high serum CD5L levels.

Our findings with CD5L-deficient mice revealed increased recruitment of monocytes, activated macrophages and neutrophils into blood characteristic of human RA [69–71]. The analogous monocytes recruitment to blood occurred with a several days delay in WT CIA mice coinciding with a drop of serum CD5L levels. This heightened inflammatory state in CD5L^-^ mice was corroborated by elevated serum levels of inflammatory cytokines IL-1β, IL-17 and IFN-γ which drive synovitis and joint damage by further attracting myeloid cells [72–75]. Notably, the increase of proinflammatory cytokines in CD5L^−^ mice converged with the increase of CD5L in WT mice, which could have dampened the cytokine release offering explanation to why it did not occur in WT mice. Together, the results of animal arthritis model and human cultured cells strongly suggest that CD5L does not promote inflammation in RA but rather serves as a protective factor. Our study also raises an interesting possibility of whether CD5L levels can pinpoint and control the developing pre-clinical phase of RA, prior to the system is overwhelmed with aggressive tissue-infiltrating macrophages accompanied by onset of inflammation.

A permanent CD5L deficiency found in human RA synovia suggests that absent CD5L compromises the system’s defense against inflammation/disease development. This implies that a) CD5L expression associates with inflammation resolving, efferocytosis macrophages profile; and b) cessation of CD5L production in monocytes/macrophages occurred prior to entering RA joints. Investigating CD14^+^ cells of RA patients, we demonstrated that serum CD5L levels and CD5L production in CD14^+^ cells resulted in a congruent anti-inflammatory evolution in monocytes. Structural integrity of CD5L with the joint chain protein linking the IgM pentamer complex [76] could explain the coincidence of serum CD5L with boosted expression of complement proteins including C1q subunits C1QA, C1QB, FCGRs and TAM/GAS6 axis.

In this study, we connected CD5L production with the basic cellular processes in human CD14^+^ cells. Production of CD5L required inhibition of histone deacetylases, the enzymes with immediate control of gene transcription. It appeared to be specifically dependent on this molecular mechanism, as neither of the pro-inflammatory stimulants, LPS/ConA or FCGR-activating IgG, alone was sufficient to increase CD5L production. Notably, we demonstrated that human leukocytes treated with HDACi downregulated production of IFN-γ, which bound together production of CD5L, IL-10 and accumulation of CD16^+^ monocyte subsets. In mice, this partnership of histone deacetylases with IFN-γ in regulation of monocyte phenotype has been shown dependent on HDAC3 and significantly affected about 50% of cytokine profile in mouse monocytes [77].

Inflammation recruits HDAC to repress macrophage genes [78]. This could be translated in upregulation of CD5L with HDAC inhibition contrasting with the view on inflammation as a strong HDAC inhibitor at early phase of arthritis. The pathogenic role of histone acetylases in arthritis has been demonstrated in mouse models while HDACi delayed arthritis development and alleviated its severity [43, 79–81], although the studies prioritized the effect of HDACi on synovial fibroblasts rather than macrophages. Our study proposes a CD5L-dependent link between the tissue trafficking of FCGR3A/CD16^+^ monocytes and development of skeletal damage.

In conclusion, this study demonstrated that the anti-inflammatory and inflammation resolving properties of CD5L are executed through induction of efferocytosis profile in monocytes. In human RA, high serum levels of CD5L harbor a potential danger of uncontrolled invasion of non-classical monocytes into synovial tissue causing joint damage. The complex role of CD5L in inflammation and disease outcomes requires careful consideration of disease phase and experimental context.

## Author contributions

D.B., E.C., M.I.B. and A.M.C. designed the experiments; M.I.B. contributed with patient material; D.B., D.S., M.C.E., E.C., M.C.S., R.F.S and L.O. performed the experiments; D.B., D.S., M.C.E., V.C., E.C., L.O., M.I.B. and A.M.C. analyzed biological data; V.L. analyzed radiographic material; D.B., D.S., M.C.E., V.C., and L.O. generated the figures; D.B., M.I.B., D.S., V.C., and A.M.C. wrote the paper; D.B., L.O., M.I.B. and A.M.C. reviewed and edited the paper. All authors read and approved the manuscript.

## Funding

This work was funded by National Funds through FCT – Fundação para a Ciência e a Tecnologia, I.P., under the project UID/4293/2025, and by grants from the Swedish Association against Rheumatism (MB, R-860371, R-994505, R-1013204), the King Gustaf V:s 80-year Foundation (MB, FAI-2020-0653, FAI-2022-0882), the Regional agreement on medical training and clinical research in the Western Götaland county (MB, ALFGBG-965623, ALFGBG-1006725), the University of Gothenburg. The funding sources have no role in study design; in the collection, analysis, and interpretation of data; in the writing of the report; and in the decision to submit the paper for publication.

The funding sources have no role in study design; in the collection, analysis, and interpretation of data; in the writing of the report; and in the decision to submit the paper for publication.

## Acknowledgments

We are thankful to the research nurses Anneli Lund and Marie-Louise Andersson, Rheumatology Clinic, Sahlgrenska University Hospital, for assistance with blood sampling. We thank all RA patients participated in this study. We thank the support of the i3S platforms Translational Cytometry and Animal Facility and Dr. Kristina Forslind for the supervision of radiographic analysis.

## Figure legends

**Supplementary Fig. S1: Gating strategy used to identify leukocyte populations in the blood of WT and CD5L⁻ mice.** Cells were identified as T cells (CD3⁺), B cells (CD19⁺), NK cells (NK1.1⁺CD3⁻), eosinophils (SSC^high^Siglec-F⁺CD11b⁺), neutrophils (Siglec-F⁻Ly6G⁺CD11b⁺), monocytes (Siglec-F⁻Ly6G⁻F4/80⁺CD11b⁺), classical/ inflammatory monocytes (Siglec-F⁻Ly6G⁻F4/80⁺CD11b⁺Ly6C⁺), and patrolling/non-classical monocytes (Siglec-F⁻Ly6G⁻F4/80⁺ CD11b⁺Ly6C⁻).

**Supplementary Fig. S2: Weight and joint histology of WT and CD5L⁻ mice in the CIA model.** Mice were immunized with chicken collagen on day 0 followed by a boost administration of the antigen on day 21. **A**, Weigh at baseline (day 0) in WT and CD5L⁻ mice cohort included in the experiments (left) and weight variation upon arthritis induction (right). Graphical representation of median and 25^th^ – 75^th^ quartiles (left) or mean with SEM (right), n= 36 (WT) and n= 39 (CD5L⁻) mice per group, pooled from 5 independent experiments. **B**, representative joint tissue sections of WT and CD5L⁻ mice recovered at day 56 post CIA induction, stained with hematoxylin and eosin. Bone (B), bone marrow (BM) and joint space (JS) are indicated. Presence of inflammatory infiltrate is indicated by black arrows in the CD5L⁻ tissue section (right).

**Supplementary Fig. S3: Dynamics of leukocyte populations and anti-collagen antibodies in the blood of WT and CD5L⁻ mice in the CIA model.** Mice were immunized with chicken collagen on day 0 followed by a boost administration of the antigen on day 21. **A**, The frequency of the indicated populations was analyzed in different time points after immunization in the blood of WT and CD5L⁻ mice by flow cytometry (frequencies within total live cells). Markers used for phenotyping: T cells (CD3^+^), B cells (CD19^+^), neutrophils (Siglec-F⁻Ly6G^+^CD11b^+^), NK cells (NK1.1^+^CD3⁻) and eosinophils (SSC^hi^Siglec-F^+^CD11b^+^). Data shown are mean with SEM of n = 10 (WT) and n= 10 (CD5L⁻) mice pooled from 2 independent experiments. Statistical comparisons were made using Šídák’s multiple comparisons test. **B**, Anti-collagen IgGs were measured in the sera of WT and CD5L⁻ mice by ELISA in the indicated timepoints following immunization. Graphical representation of median and 25^th^ – 75^th^ quartiles. Statistical significance is indicated as \**p* < 0.05; \*\****p*** < 0.01.

**Supplementary Fig. S4: A**, Gating strategy and representative dot plot of CD11b^+^CD4^-^ monocyte population, in flow cytometry. **B,** Box plot of monocyte subtype frequency in CD11b^+^CD4^-^ cells stimulated as indicated, by flow cytometry. **C,** Protein levels of CD5L in PBMC cultures measured by ELISA. **D,** Protein levels of IFN-γ in PBMC cultures measured by ELISA. LPS/ConA, lipopolysaccharide/concanavalin A; HDACi, histone deacetylase inhibitor; LynAct, Lyn activator.

**Supplementary Fig. S5: A**, Protein levels of cytokines in supernatants of CD14^+^ cells of RA patients with high (above twice of the detection level, 0.6 pg/ml, n=10) and low levels of CD5L (n=25) dichotomized by median. **B,** Biological processes enriched in transcriptome of CD5L-producing CD14^+^ cells. Enrichment analysis is done in GSEA. **C,** Heat map of gene expression (log_2_ fold change, FC) in regression to CD5L levels in serum, in supernatants of CD14^+^ cells, and to vdH-Sharp score. Analysis was done by DESeq2 method. P values were calculated using DESeq2; nominal P values are indicated as \**p* < 0.05; \*\****p*** < 0.01; \*\*\****p*** < 0.001.

## Notes

### Competing Interest Statement

The authors have declared no competing interest.

## References

1. Miyazaki, T., et al., Increased susceptibility of thymocytes to apoptosis in mice lacking AIM, a novel murine macrophage-derived soluble factor belonging to the scavenger receptor cysteine-rich domain superfamily. J. Exp. Med., 1999. 189(2): p. 413–422.

2. Sarrias, M.R., et al., Biochemical characterization of recombinant and circulating human Spalpha. Tissue Antigens, 2004. 63(4): p. 335–344.

3. Gebe, J.A., et al., Molecular cloning, mapping to human chromosome 1 q21-q23, and cell binding characteristics of Spalpha, a new member of the scavenger receptor cysteine-rich (SRCR) family of proteins. J. Biol. Chem., 1997. 272(10): p. 6151–6158.

4. Sarrias, M.R., et al., A role for human Sp alpha as a pattern recognition receptor. J. Biol. Chem., 2005. 280(42): p. 35391–35398.

5. Martínez, V.G., et al., The macrophage soluble receptor AIM/Api6/CD5L displays a broad pathogen recognition spectrum and is involved in early response to microbial aggression. Cell. Mol. Immunol., 2014. 11(4): p. 343–354.

6. Arai, S., et al., Apoptosis inhibitor of macrophage protein enhances intraluminal debris clearance and ameliorates acute kidney injury in mice. Nat Med, 2016. 22(2): p. 183–93.

7. Maehara, N., et al., AIM/CD5L attenuates DAMPs in the injured brain and thereby ameliorates ischemic stroke. Cell Rep, 2021. 36(11): p. 109693.

8. Kurokawa, J., et al., Macrophage-derived AIM is endocytosed into adipocytes and decreases lipid droplets via inhibition of fatty acid synthase activity. Cell Metab, 2010. 11(6): p. 479–92.

9. Sanchez-Moral, L., et al., Multifaceted Roles of CD5L in Infectious and Sterile Inflammation. Int J Mol Sci, 2021. 22(8).

10. Oliveira, L., et al., CD5L as a promising biological therapeutic for treating sepsis. Nat Commun, 2024. 15(1): p. 4119.

11. Wang, C., et al., CD5L/AIM Regulates Lipid Biosynthesis and Restrains Th17 Cell Pathogenicity. Cell, 2015. 163(6): p. 1413–27.

12. Guo, Y., M. Zhu, and R. Shen, CD5L Deficiency Protects Mice Against Bleomycin-Induced Pulmonary Fibrosis. Front Biosci (Landmark Ed), 2023. 28(9): p. 209.

13. Li, M., et al., CD5L deficiency attenuate acetaminophen-induced liver damage in mice via regulation of JNK and ERK signaling pathway. Cell Death Discov, 2021. 7(1): p. 342.

14. Cardoso, M.S., et al., CD5L is upregulated upon infection with Mycobacterium tuberculosis with no effect on disease progression. Immunology, 2024.

15. Kim, W.K., et al., Glycoproteomic analysis of plasma from patients with atopic dermatitis: CD5L and ApoE as potential biomarkers. Exp Mol Med, 2008. 40(6): p. 677–85.

16. Lai, X., et al., Elevation of serum CD5L concentration is correlated with disease activity in patients with systemic lupus erythematosus. Int Immunopharmacol, 2018. 63: p. 311–316.

17. Gangadharan, B., et al., Novel serum biomarker candidates for liver fibrosis in hepatitis C patients. Clin Chem, 2007. 53(10): p. 1792–9.

18. Riedel, B.C., et al., Uncovering Biologically Coherent Peripheral Signatures of Health and Risk for Alzheimer’s Disease in the Aging Brain. Front Aging Neurosci, 2018. 10: p. 390.

19. Yu, H.R., et al., A unique plasma proteomic profiling with imbalanced fibrinogen cascade in patients with Kawasaki disease. Pediatr Allergy Immunol, 2009. 20(7): p. 699–707.

20. Wu, X., et al., Apoptosis inhibitor of macrophage/CD5L is associated with disease activity in rheumatoid arthritis. Clin Exp Rheumatol, 2021. 39(1): p. 58–65.

21. Almutairi, K.B., et al., The Prevalence of Rheumatoid Arthritis: A Systematic Review of Population-based Studies. J Rheumatol, 2021. 48(5): p. 669–676.

22. Isaacs, J.D., The changing face of rheumatoid arthritis: sustained remission for all? Nat Rev Immunol, 2010. 10(8): p. 605–11.

23. Gravallese, E.M. and G.S. Firestein, Rheumatoid Arthritis – Common Origins, Divergent Mechanisms. N Engl J Med, 2023. 388(6): p. 529–542.

24. Saeki, N. and Y. Imai, Crosstalk between synovial macrophages and fibroblasts in rheumatoid arthritis. Histol Histopathol, 2023. 38(11): p. 1231–1238.

25. Murray, P.J., et al., Macrophage activation and polarization: nomenclature and experimental guidelines. Immunity, 2014. 41(1): p. 14–20.

26. Salnikova, D.I., et al., Target Role of Monocytes as Key Cells of Innate Immunity in Rheumatoid Arthritis. Diseases, 2024. 12(5).

27. Iwamoto, T., et al., Molecular aspects of rheumatoid arthritis: chemokines in the joints of patients. FEBS J, 2008. 275(18): p. 4448–55.

28. Rana, A.K., et al., Monocytes in rheumatoid arthritis: Circulating precursors of macrophages and osteoclasts and, their heterogeneity and plasticity role in RA pathogenesis. Int Immunopharmacol, 2018. 65: p. 348–359.

29. Chomarat, P., et al., IL-6 switches the differentiation of monocytes from dendritic cells to macrophages. Nat Immunol, 2000. 1(6): p. 510–4.

30. Yang, X., Y. Chang, and W. Wei, Emerging role of targeting macrophages in rheumatoid arthritis: Focus on polarization, metabolism and apoptosis. Cell Prolif, 2020. 53(7): p. e12854.

31. Cutolo, M., et al., The Role of M1/M2 Macrophage Polarization in Rheumatoid Arthritis Synovitis. Front Immunol, 2022. 13: p. 867260.

32. Ospelt, C., Synovial fibroblasts in 2017. RMD Open, 2017. 3(2): p. e000471.

33. Fangradt, M., et al., Human monocytes and macrophages differ in their mechanisms of adaptation to hypoxia. Arthritis Res Ther, 2012. 14(4): p. R181.

34. Tanaka, M., et al., Activation of Fc gamma RI on monocytes triggers differentiation into immature dendritic cells that induce autoreactive T cell responses. J Immunol, 2009. 183(4): p. 2349–55.

35. Alivernini, S., et al., Distinct synovial tissue macrophage subsets regulate inflammation and remission in rheumatoid arthritis. Nat Med, 2020. 26(8): p. 1295–1306.

36. Kuo, D., et al., HBEGF(+) macrophages in rheumatoid arthritis induce fibroblast invasiveness. Sci Transl Med, 2019. 11(491).

37. Culemann, S., et al., Locally renewing resident synovial macrophages provide a protective barrier for the joint. Nature, 2019. 572(7771): p. 670–675.

38. Faas, M., et al., IL-33-induced metabolic reprogramming controls the differentiation of alternatively activated macrophages and the resolution of inflammation. Immunity, 2021. 54(11): p. 2531–2546 e5.

39. Cai, B., et al., MerTK signaling in macrophages promotes the synthesis of inflammation resolution mediators by suppressing CaMKII activity. Sci Signal, 2018. 11(549).

40. Klein, K., C. Ospelt, and S. Gay, Epigenetic contributions in the development of rheumatoid arthritis. Arthritis Res Ther, 2012. 14(6): p. 227.

41. Yang, C., et al., Epigenetic Regulation in the Pathogenesis of Rheumatoid Arthritis. Front Immunol, 2022. 13: p. 859400.

42. Li, Y., et al., Reduced Activity of HDAC3 and Increased Acetylation of Histones H3 in Peripheral Blood Mononuclear Cells of Patients with Rheumatoid Arthritis. J Immunol Res, 2018. 2018: p. 7313515.

43. Angiolilli, C., et al., Histone deacetylase 3 regulates the inflammatory gene expression programme of rheumatoid arthritis fibroblast-like synoviocytes. Ann Rheum Dis, 2017. 76(1): p. 277–285.

44. Nogueira, E., et al., Enhancing Methotrexate Tolerance with Folate Tagged Liposomes in Arthritic Mice. Journal of Biomedical Nanotechnology, 2015. 11(12): p. 2243–2252.

45. Strait, R.T., S. Thornton, and F.D. Finkelman, Cgamma1 Deficiency Exacerbates Collagen-Induced Arthritis. Arthritis Rheumatol, 2016. 68(7): p. 1780–7.

46. Pincus, T., R.H. Brooks, and L.F. Callahan, A proposed 30-45 minute 4 page standard protocol to evaluate rheumatoid arthritis (SPERA) that includes measures of inflammatory activity, joint damage, and longterm outcomes. J Rheumatol, 1999. 26(2): p. 473–80.

47. Aletaha, D., J. Smolen, and M.M. Ward, Measuring function in rheumatoid arthritis: Identifying reversible and irreversible components. Arthritis Rheum, 2006. 54(9): p. 2784–92.

48. van der Heijde, D.M., et al., Biannual radiographic assessments of hands and feet in a three-year prospective followup of patients with early rheumatoid arthritis. Arthritis Rheum, 1992. 35(1): p. 26–34.

49. Helle, M., L. Boeije, and L.A. Aarden, Functional discrimination between interleukin 6 and interleukin 1. Eur J Immunol, 1988. 18(10): p. 1535–40.

50. Zhang, F., et al., Deconstruction of rheumatoid arthritis synovium defines inflammatory subtypes. Nature, 2023. 623(7987): p. 616–624.

51. Blanco-Carmona, E., Generating publication ready visualizations for Single Cell transcriptomics using SCpubr. bioRxiv, 2022: p. 2022.02.28.482303.

52. Marsh, S.E., scCustomize: Custom Visualizations & Functions for Streamlined Analyses of Single Cell Sequencing. 2021, GitHub.

53. Hulsebus, H.J., et al., Complement Component C1q Programs a Pro-Efferocytic Phenotype while Limiting TNFalpha Production in Primary Mouse and Human Macrophages. Front Immunol, 2016. 7: p. 230.

54. Binvignat, M., et al., Single-cell RNA-Seq analysis reveals cell subsets and gene signatures associated with rheumatoid arthritis disease activity. JCI Insight, 2024. 9(16).

55. Mulder, K., et al., Cross-tissue single-cell landscape of human monocytes and macrophages in health and disease. Immunity, 2021. 54(8): p. 1883–1900 e5.

56. Wang, W., et al., Identification of hypoxic macrophages in glioblastoma with therapeutic potential for vasculature normalization. Cancer Cell, 2024. 42(5): p. 815–832 e12.

57. Yasuda, K., et al., AIM/CD5L ameliorates autoimmune arthritis by promoting removal of inflammatory DAMPs at the lesions. J Autoimmun, 2024. 142: p. 103149.

58. Sakthiswary, R., et al., Cartilage oligomeric matrix protein (COMP) in rheumatoid arthritis and its correlation with sonographic knee cartilage thickness and disease activity. Clin Rheumatol, 2017. 36(12): p. 2683–2688.

59. Lindqvist, E., et al., Prognostic laboratory markers of joint damage in rheumatoid arthritis. Ann Rheum Dis, 2005. 64(2): p. 196–201.

60. Gao, L., et al., Receptor tyrosine kinases Tyro3, Axl, and Mertk differentially contribute to antibody-induced arthritis. Cell Commun Signal, 2023. 21(1): p. 195.

61. Geven, E.J., et al., S100A8/A9, a potent serum and molecular imaging biomarker for synovial inflammation and joint destruction in seronegative experimental arthritis. Arthritis Res Ther, 2016. 18(1): p. 247.

62. Kang, K., et al., Interferon-gamma Represses M2 Gene Expression in Human Macrophages by Disassembling Enhancers Bound by the Transcription Factor MAF. Immunity, 2017. 47(2): p. 235–250 e4.

63. Misharin, A.V., et al., Nonclassical Ly6C(-) monocytes drive the development of inflammatory arthritis in mice. Cell Rep, 2014. 9(2): p. 591–604.

64. Puchner, A., et al., Non-classical monocytes as mediators of tissue destruction in arthritis. Ann Rheum Dis, 2018. 77(10): p. 1490–1497.

65. Tanaka, M., S.S. Dykes, and D.W. Siemann, Inhibition of the Axl pathway impairs breast and prostate cancer metastasis to the bones and bone remodeling. Clin Exp Metastasis, 2021. 38(3): p. 321–335.

66. Nassar, M., et al., GAS6 is a key homeostatic immunological regulator of host-commensal interactions in the oral mucosa. Proc Natl Acad Sci U S A, 2017. 114(3): p. E337–E346.

67. Villanueva-Martin, G., et al., Non-classical circulating monocytes expressing high levels of microsomal prostaglandin E2 synthase-1 tag an aberrant IFN-response in systemic sclerosis. J Autoimmun, 2023. 140: p. 103097.

68. Sanjurjo, L., et al., CD5L Promotes M2 Macrophage Polarization through Autophagy-Mediated Upregulation of ID3. Front Immunol, 2018. 9: p. 480.

69. Cascao, R., et al., Neutrophils in rheumatoid arthritis: More than simple final effectors. Autoimmun Rev, 2010. 9(8): p. 531–5.

70. Liote, F., et al., Blood monocyte activation in rheumatoid arthritis: increased monocyte adhesiveness, integrin expression, and cytokine release. Clin Exp Immunol, 1996. 106(1): p. 13–9.

71. Laria, A., et al., The macrophages in rheumatic diseases. J Inflamm Res, 2016. 9: p. 1–11.

72. Kuwabara, T., et al., The Role of IL-17 and Related Cytokines in Inflammatory Autoimmune Diseases. Mediators Inflamm, 2017. 2017: p. 3908061.

73. Palmblad, K., et al., Dynamics of early synovial cytokine expression in rodent collagen-induced arthritis: a therapeutic study using a macrophage-deactivating compound. Am J Pathol, 2001. 158(2): p. 491–500.

74. Srirangan, S. and E.H. Choy, The role of interleukin 6 in the pathophysiology of rheumatoid arthritis. Ther Adv Musculoskelet Dis, 2010. 2(5): p. 247–56.

75. Kato, M., New insights into IFN-gamma in rheumatoid arthritis: role in the era of JAK inhibitors. Immunol Med, 2020. 43(2): p. 72–78.

76. Oskam, N., et al., CD5L is a canonical component of circulatory IgM. Proc Natl Acad Sci U S A, 2023. 120(50): p. e2311265120.

77. Chen, X., et al., Requirement for the histone deacetylase Hdac3 for the inflammatory gene expression program in macrophages. Proc Natl Acad Sci U S A, 2012. 109(42): p. E2865–74.

78. Marques, O., et al., Inflammation-driven NF-kappaB signaling represses ferroportin transcription in macrophages via HDAC1 and HDAC3. Blood, 2025. 145(8): p. 866–880.

79. Joosten, L.A., et al., Inhibition of HDAC activity by ITF2357 ameliorates joint inflammation and prevents cartilage and bone destruction in experimental arthritis. Mol Med, 2011. 17(5-6): p. 391–6.

80. Lin, H.S., et al., Anti-rheumatic activities of histone deacetylase (HDAC) inhibitors in vivo in collagen-induced arthritis in rodents. Br J Pharmacol, 2007. 150(7): p. 862–72.

81. Grabiec, A.M., et al., Histone deacetylase inhibitors suppress rheumatoid arthritis fibroblast-like synoviocyte and macrophage IL-6 production by accelerating mRNA decay. Ann Rheum Dis, 2012. 71(3): p. 424–31.

