## Supplementary Figures for "CD5L insufficiency exacerbates skeletal joint damage in rheumatoid arthritis"

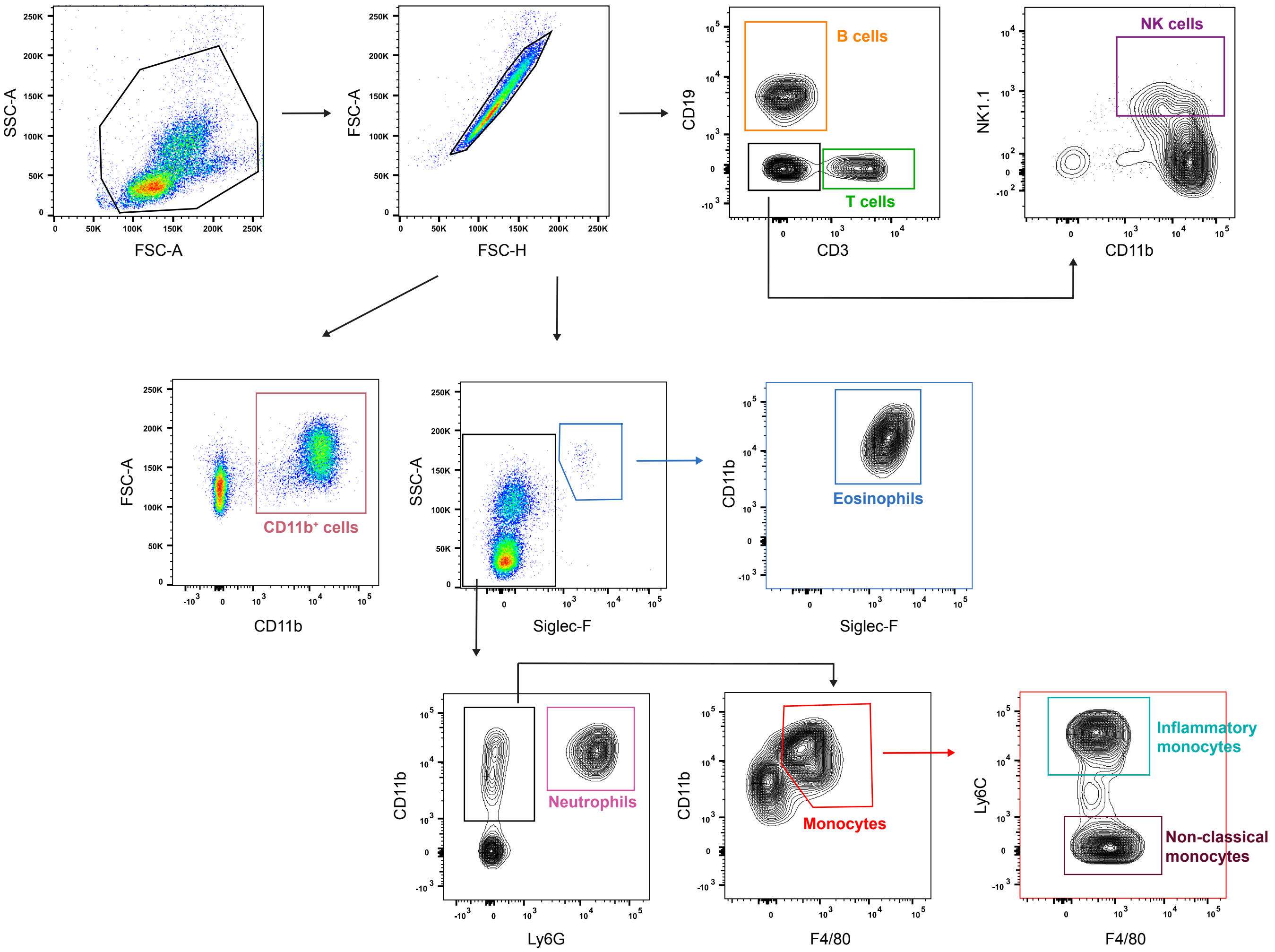


**Figure S1.** *Gating strategy used to identify leukocyte populations in the blood of WT and CD5L⁻ mice.* Cells were identified as T cells ($CD3^+$), B cells ($CD19^+$), NK cells ($NK1.1^+CD3^-$), eosinophils ($SSC^{high}$Siglec-$F^+CD11b^+$), neutrophils (Siglec-$F^-Ly6G^+CD11b^+$), monocytes (Siglec-$F^-Ly6G^-F4/80^+CD11b^+$), classical/inflammatory monocytes (Siglec-$F^-Ly6G^-F4/80^+CD11b^+Ly6C^+$), and patrolling/non-classical monocytes (Siglec-$F^-Ly6G^-F4/80^+$ $CD11b^+Ly6C^-$).

### Supplementary Figure 2

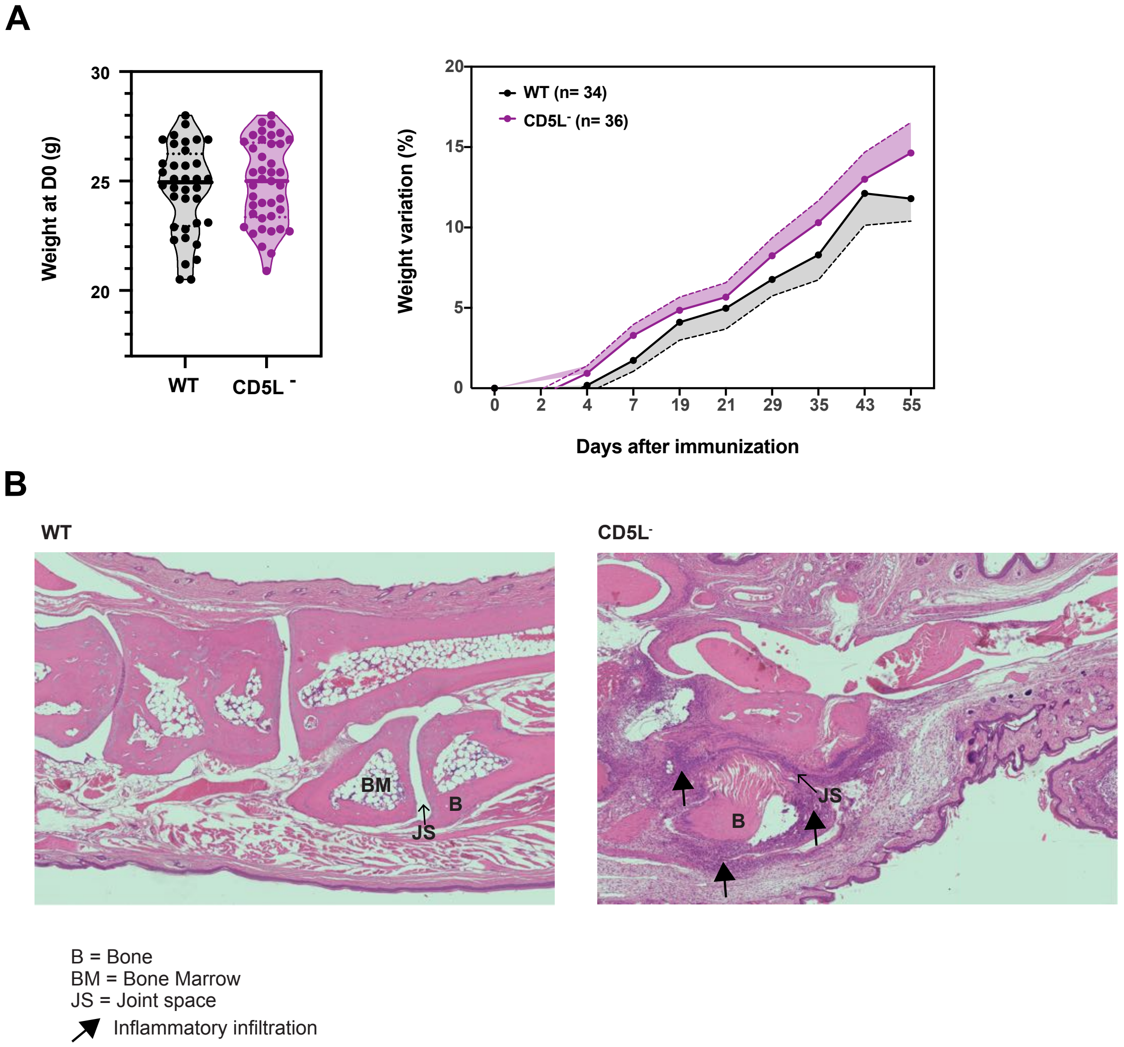


**Figure S2 – Weight and joint histology of WT and CD5L- mice in the CIA model.** Mice were immunized with chicken collagen on day 0 followed by a boost administration of the antigen on day 21. **A**) Weigh at baseline (day 0) in WT and CD5L- mice cohort included in the experiments (left) and weight variation upon arthritis induction (right). Graphical representation of median and $25^{th}$ - $75^{th}$ quartiles (left) or mean with SEM (right), n= 36 (WT) and n= 39 (CD5L-) mice per group, pooled from 5 independent experiments. **B)** representative joint tissue sections of WT and CD5L- mice recovered at day 56 post CIA induction, stained with hematoxylin and eosin. Bone (B), bone marrow (BM) and joint space (JS) are indicated. Presence of inflammatory infiltrate is indicated by black arrows in the CD5L- tissue section (right).

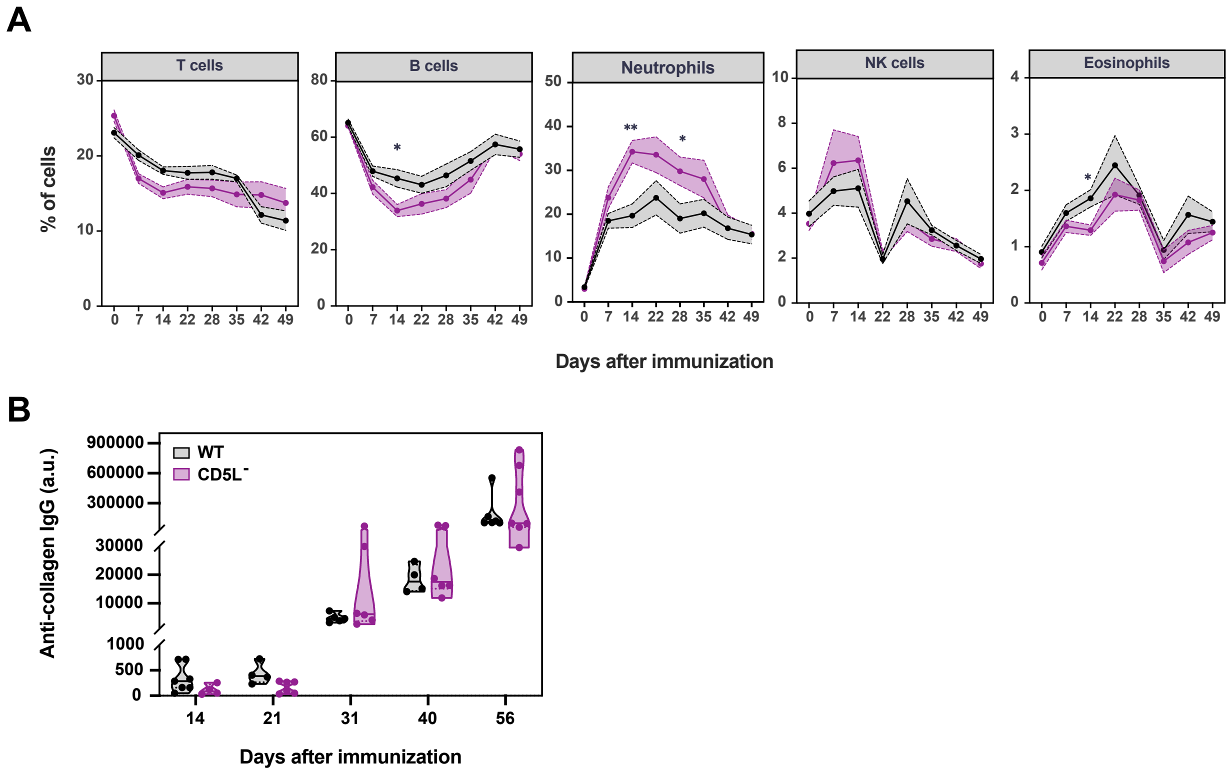


**Figure S3 – Dynamics of leukocyte populations and anti-collagen antibodies in the blood of WT and CD5L$^-$ mice in the CIA model.** Mice were immunized with chicken collagen on day 0 followed by a boost administration of the antigen on day 21. **A)** The frequency of the indicated populations was analyzed in different time points after immunization in the blood of WT and CD5L$^-$ mice by flow cytometry (frequencies within total live cells). Markers used for phenotyping: T cells (CD3$^+$), B cells (CD19$^+$), neutrophils (Siglec-F$^-$Ly6G$^+$CD11b$^+$), NK cells (NK1.1$^+$ CD3$^-$) and eosinophils (SSC$^{high}$ Siglec-F$^+$ CD11b$^+$). Data shown are mean with SEM of n = 10 (WT) and n= 10 (CD5L$^-$) mice pooled from 2 independent experiments. Statistical comparisons were made using Šídák's multiple comparisons test. **B)** Anti-collagen IgGs were measured in the sera of WT and CD5L$^-$ mice by ELISA in the indicated timepoints following immunization. Graphical representation of median and 25$^{th}$ - 75$^{th}$ quartiles. *$P$ < 0.05, **$P$<0.01.

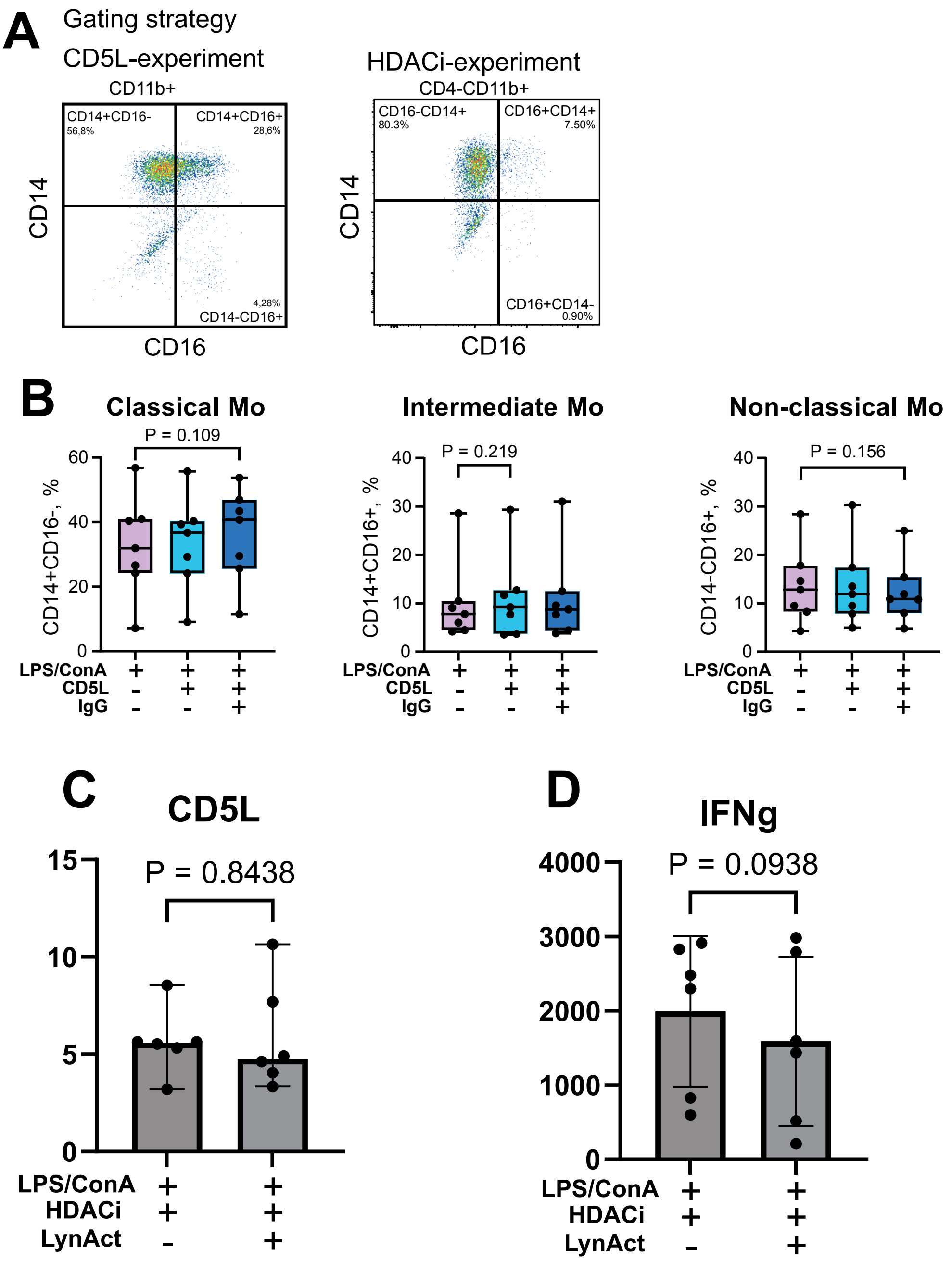


**Supplementary Figure S4: A,** Gating strategy and representative dot plot of CD11b$^{+}$CD4$^{-}$ monocyte population, in flow cytometry. **B,** Box plot of monocyte subtype frequency in CD11b$^{+}$CD4$^{-}$ cells stimulated as indicated, by flow cytometry. **C,** Protein levels of CD5L in PBMC cultures measured by ELISA. **D,** Protein levels of IFNg in PBMC cultures measured by ELISA. LPS/ConA, lipopolysaccharide/concanavalin A; HDACi, histone deacetylase inhibitor, LynAct, Lyn activator.

**Supplementary figure 5**

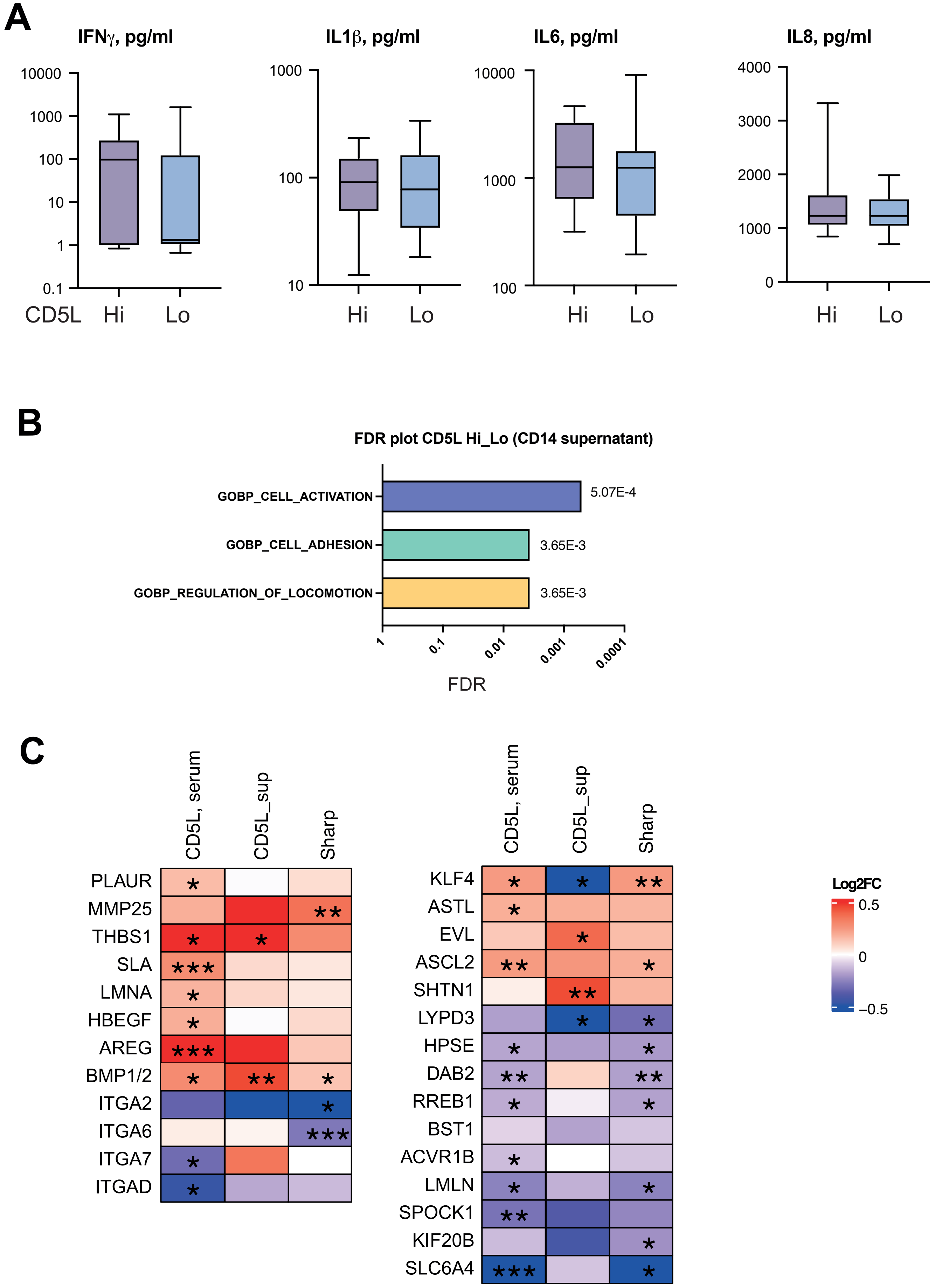


**Supplementary Figure S5: A,** Protein levels of cytokines in supernatants of CD14+ cells of RA patients with high (above twice of the detection level, 0.6 pg/ml, n=10) and low levels of CD5L (n=25) dichotomized by median. **B,** Biological processes enriched in transcriptome of CD5L-producing CD14$^{+}$ cells. Enrichment analysis is done in GSEA. **C,** Heat map of gene expression ($log_2$ fold change, FC) in regression to CD5L levels in serum, in supernatants of CD14$^{+}$ cells, and to vdH-Sharp score. Analysis was done by DESeq2 method. P values were calculated using DESeq2; nominal P values are indicated as *p < 0.05; **p < 0.01; ***p < 0.001.
